# Non-human primates satisfy utility maximization in compliance with the continuity axiom of Expected Utility Theory

**DOI:** 10.1101/2020.02.18.953950

**Authors:** Simone Ferrari-Toniolo, Philipe M. Bujold, Fabian Grabenhorst, Raymundo Báez-Mendoza, Wolfram Schultz

## Abstract

Expected Utility Theory (EUT), the first axiomatic theory of risky choice, describes choices as a utility maximization process: decision makers assign a subjective value (utility) to each choice option and choose the one with the highest utility. The continuity axiom, central to EUT and its modifications, is a necessary and sufficient condition for the definition of numerical utilities. The axiom requires decision makers to be indifferent between a gamble and a specific probabilistic combination of a more preferred and a less preferred gamble. While previous studies demonstrated that monkeys choose according to combinations of objective reward magnitude and probability, a concept-driven experimental approach for assessing the axiomatically defined conditions for maximizing subjective utility by animals is missing. We experimentally tested the continuity axiom for a broad class of gamble types in four male rhesus macaque monkeys, showing that their choice behavior complied with the existence of a numerical utility measure as defined by the economic theory. We used the numerical quantity specified in the continuity axiom to characterize subjective preferences in a magnitude-probability space. This mapping highlighted a trade-off relation between reward magnitudes and probabilities, compatible with the existence of a utility function underlying subjective value computation. These results support the existence of a numerical utility function able to describe choices, allowing for the investigation of the neuronal substrates responsible for coding such rigorously defined quantity.

**SIGNIFICANCE STATEMENT:** A common assumption of several economic choice theories is that decisions result from the comparison of subjectively assigned values (utilities). This study demonstrated the compliance of monkey behavior with the continuity axiom of Expected Utility Theory, implying a subjective magnitude-probability trade-off relation which supports the existence of numerical subjective utility directly linked to the theoretical economic framework. We determined a numerical utility measure able to describe choices, which can serve as a correlate for the neuronal activity in the quest for brain structures and mechanisms guiding decisions.

## INTRODUCTION

Expected Utility Theory (EUT), the first axiomatic theory of risky choice (von Neumann and Morgenstern, 1944), demonstrates that if a subject’s behavior followed a simple set of rules, or axioms, their choices could be described by utility maximization (Methods section: *Expected Utility theorem*). The utility concept is used by economists to mathematically define subjective values, and its maximization is a general and basic process determining the subject’s survival. The core of EUT remained central to the generalized Expected Utility theories developed later, most notably Prospect Theory (Kahneman and Tversky, 1979; Hey and Orme, 1994; Starmer, 2000), which overcame the limits of EUT by introducing the concepts of probability weighting and reference point.

The continuity axiom of EUT, given three subjectively ranked gambles, requires the existence of an indifference point (IP) between the intermediate gamble and a probabilistic combination of the other two. This ensures that no option is considered infinitely better than any other option, making it possible to define a finite, numerical subjective value for each gamble (Jehle and Reny, 2001; Chateauneuf et al., 2008; Johnson and Ratcliff, 2014). Continuity is common to a broad spectrum of risky and riskless choice theories (Samuelson, 1948; Weber and Camerer, 1987), emerging as a fundamental construct in all economic schemes based on value computation.

Compliance with the continuity axiom can be experimentally tested by verifying the existence of indifference between the intermediate gamble and one of the probabilistic combinations. Violations according to the mathematical definition cannot be experimentally tested, as they would require infinitesimally small probability steps and thus an infinite number of tests. Nevertheless, given the biological limits of perceiving small probability differences (Tversky, 1969), tests within the finite range of noticeable probability differences could be interpreted as being biologically plausible, and finding violations in such limited-range tests could be considered relevant.

Monkeys are the evolutionary closest species to humans that allow to investigate, at the single-cell level and over multiple repetitions, the neuronal correlates of economic choice between options with finely varying reward magnitude and probability. Monkeys prefer gambles with larger EVs to those with smaller EVs derived from both reward magnitude and probability (Musallam et al., 2004; Lau and Glimcher, 2005; Averbeck, 2015; Farashahi et al., 2018). These results suggested a trade-off between reward magnitude and probability, but their interpretation relies tacitly on the validity of the EUT continuity axiom. The axiom would formally distinguish between circumstantial and systematic trade-offs but was not tested in these studies. Further, behavioral compliance with EUT axioms would provide a conceptual foundation for the meaningful neuronal coding of subjective value and formal economic utility reported previously (Platt and Glimcher, 1999; Tremblay and Schultz, 1999; Padoa-Schioppa and Assad, 2006; Kobayashi and Schultz, 2008; Lak et al., 2014; Stauffer et al., 2014).

Here, we tested whether economic choices of rhesus monkeys comply with the EUT continuity axiom, assuming no adaptation in reference point. The animals chose between a gamble and a probabilistic combination of a more preferred and a less preferred gamble. The animals revealed their preference by selecting one of the two options. Our statistical test procedure (preferences ranked with probability, the combined option going from non-preferred to preferred) identified significant compliance with the axiom. To address the general axiom definition, we tested a broad range of magnitudes and probabilities, starting with degenerate gambles (i.e. only one outcome, probability P=1.0) and advancing to gradually more complex gambles containing two or three possible outcomes.

We found that the choices of all four animals complied with the continuity axiom by expressing consistent preferences which robustly identified IPs across the tested magnitude-probability space. The animals’ behavior aligned closely with choice functions that modelled the magnitude-probability tradeoff. This result suggests the existence of an axiomatically defined utility measure for choice options. Compatible with economic theory, the pattern of subjective IPs allowed us to define a utility function that is suitable for investigating the neuronal mechanisms of risky choice.

## MATERIALS AND METHODS

### Animals and ethical approval

Four adult male macaque monkeys (*Macaca mulatta*), were used in these experiments (weight, per animal: 12.7 kg (Monkey A), 13.8 kg (Monkey B), 10.3 kg (Monkey C) and 12.5 kg (Monkey D)).

This research has been ethically reviewed, approved, regulated and supervised by the following institutions and individuals in the UK and at the University of Cambridge (UCam): the UK Home Office implementing the Animals (Scientific Procedures) Act 1986 with Amendment Regulations 2012, the local UK Home Office Inspector, the UK Animals in Science Committee (ASC), the UK National Centre for Replacement, Refinement and Reduction of Animal Experiments (NC3Rs), two UCam Animal Welfare and Ethical Review Bodies (AWERBs), the UCam Governance and Strategy Committee, the Certificate Holder of the UCam Biomedical Service (UBS), the UBS Director for Governance and Welfare, the UBS Named Veterinary Surgeon (NVS), and the UBS Named Animal Care and Welfare Officer (NACWO).

### Experimental Design and Statistical Analysis

We presented each animal with a set of two mutually exclusive and collectively exhaustive options that appeared simultaneously and at equal joystick distance in front of the animal on a computer monitor. In each option, we set the magnitude and probability of reward independently.

We tested choices by implementing the following basic concepts:

1. Increasing the reward probability in one option, while holding constant the reward magnitude in this option and the reward probability and magnitude in its alternative, leads to monotonically increasing probability of choosing this option over its alternative, as modeled by an S-shaped psychophysical choice function (Figs. 1e; 2).
2. Options are equally revealed preferred, and inferred to have equal utility for the animal, when the animal chooses them with equal probability (*P* = 0.5 each option).
3. Each option with its specific reward magnitude and probability is graphically represented at the intersection of the x-coordinate (probability) and y-coordinate (magnitude) of a two-dimensional plot (Fig. 4).
4. An option that is as revealed preferred as another option, as shown by choice indifference, is graphically represented as a two-dimensional indifference point (IP). The dots in Fig. 4b-d are IPs relative to all other color-matched dots.
5. Several IPs align as an indifference curve (IC) on which each option is as revealed preferred as any other option on that same IC, despite different magnitude-probability composition. The colored curves in Fig. 4b, d are ICs. Options on higher ICs (farther from origin) are revealed preferred to options on lower ICs, and all options on a given IC are revealed preferred to options below that IC. Options on the red and orange ICs in Fig. 4c, d are revealed preferred to options on the blue and green ICs.

**Figure 1.**
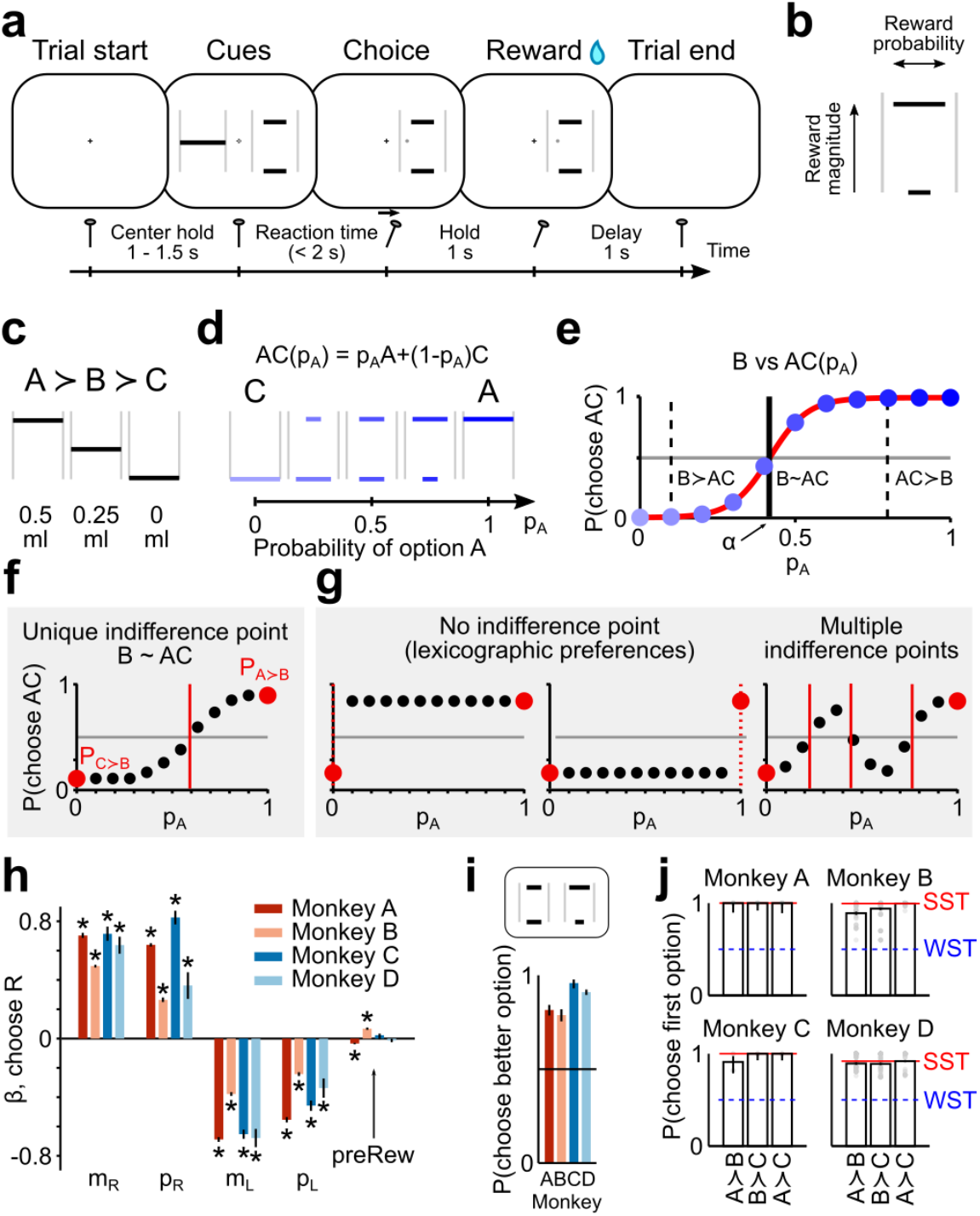
Experimental design and consistency of choice behavior. **a) Trial sequence**. Monkeys chose between two options by moving a cursor (gray dot) with a joystick to one side of the screen. After a delay, the reward corresponding to the selected cue was delivered. **b) Visual cues** indicated magnitude and probability of possible outcomes through horizontal bars’ vertical position and width, respectively. **c,d,e) Continuity axiom test**. The continuity axiom was tested through choices between a fixed gamble B and a probabilistic combination of A and C (AC). A, B and C were ordered reward magnitudes (**c**); AC was a gamble between A and C, with probabilities p_A_ and 1-p_A_ respectively (**d**); different shades of blue correspond to different p_A_ values (darker for higher p_A_). The continuity axiom implies the existence of a unique AC combination (p_A_=α) corresponding to choice indifference between the two options (B~AC, vertical line in **e**), with the existence of a p_A_ for which B≻AC and of a different p_A_ for which AC≻B (vertical dashed lines). The value of α was identified by fitting a softmax function (Eq. 2, red line) to the proportion of AC choices (blue dots). **f,g) Compliance and violation.** Choice pattern compatible with the continuity axiom (**f**) and possible axiom violations (**g**). Red dots represent the proportion of AC choices when pA=0 or 1, corresponding to the axiom’s initial requirement (A≻B and B≻C, implying P(A≻B)>0.5 and P(C≻B)<0.5). **h,i,j) Consistency of choice behavior.** The standardized beta coefficients from logistic regressions of single trials’ behavior (**h**) showed that the main choice-driving variables were reward magnitude (mR, mL) and probability (p_R_, p_L_) for all animals, both for left (L) and right (R) choices; previous trial’s chosen side (preCh_R_) and reward (preRew_R_) did not consistently explain animals’ choices (error bars: 95% CI across sessions; * *p*<0.05, one-sample *t* test, FDR corrected; no. of sessions per animal: 100 (A), 81 (B), 24 (C), 15 (D)). In choices between options with different probability of delivering the same reward magnitude, the better option was preferred on average by all animals, demonstrating compliance with FSD (**i**) (error bars: binomial 95% CI; no. of tests per animal: 28 (A), 24 (B), 15 (C), 23 (D); average no. of trials per test: 12 (A), 13 (B), 11 (C), 34 (D)). In choices between sure rewards (bars: average across all sessions; gray dots: single sessions; error bars: binomial 95% CI) animals preferred A to B, B to C and A to C (**j**), complying with both weak and strong stochastic transitivity (WST: proportion of choices of the better option >0.5 (blue dashed line); SST: proportion of A over C choices (red line) ≥ other choice proportions).

**Figure 2.**
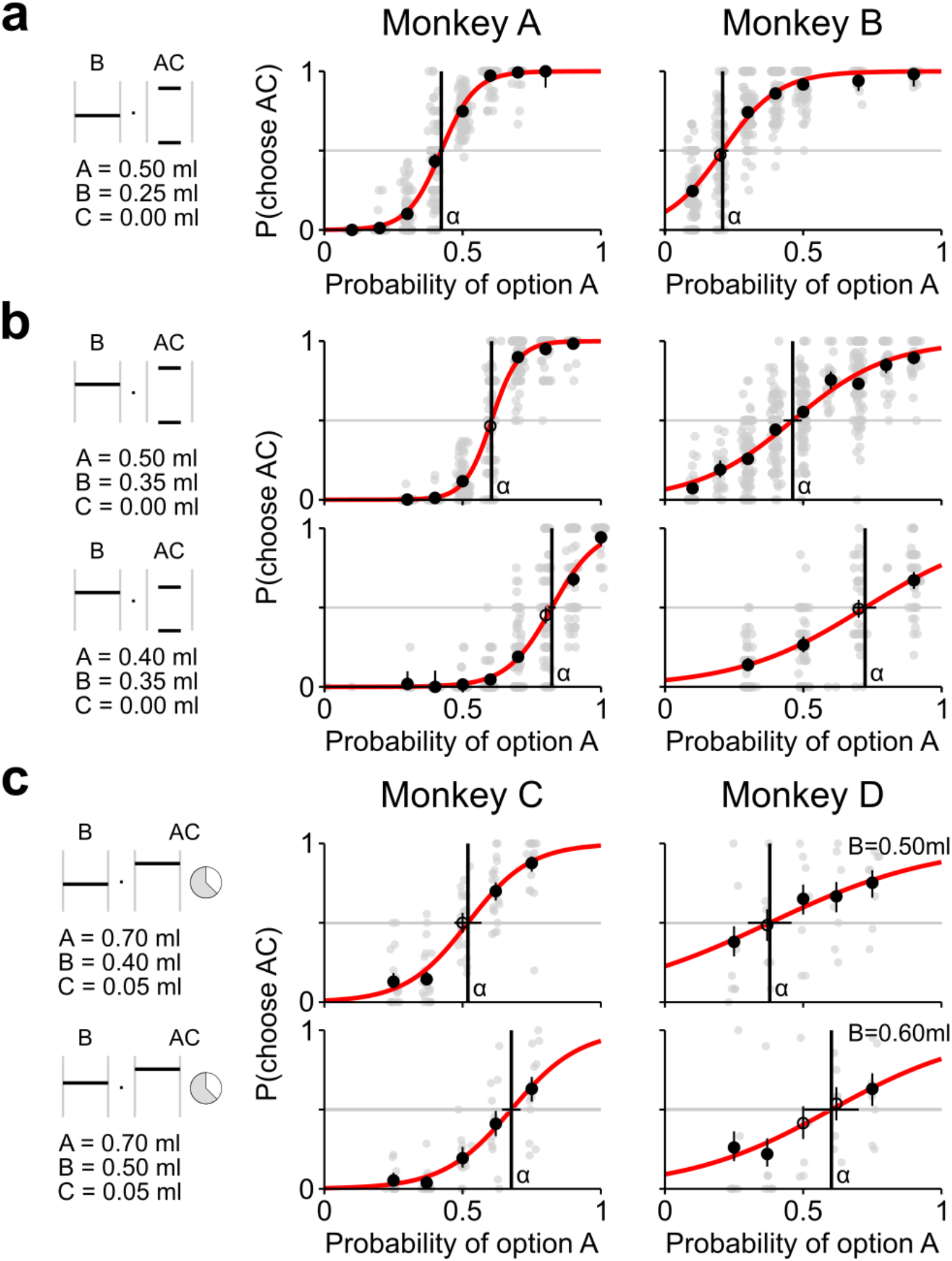
Experimental test of the continuity axiom. **a,b,c) Compliance with the continuity axiom**. The axiom was tested through choices between a gamble B and a varying AC-combined gamble (left: visual stimuli for an example choice pair with p_A_=0.5 (**a,b**) or p_A_=0.375 (c)); increasing p_A_ values resulted in gradually increasing preferences for the AC option. In each plot, gray dots represent the proportion of AC choices in single sessions, black circles the proportions across all tested sessions with vertical bars indicating the binomial 95% confidence intervals (filled circles indicate significant difference from 0.5; binomial test, *p*<0.05). The tests were repeated using different A and B values (**b**) as well as non-zero C values in a modified task (**c**). All four animals complied with the continuity axiom by showing increasing preferences for increasing probability of gamble A (rank correlation, *p*<0.05), with the AC option switching from non-preferred (p_choose AC_<0.5) to preferred (p_choose AC_>0.5) (binomial test, *p*<0.05). Each IP (α, vertical line) was computed as the p_A_ for which a data-fitted softmax function (Eq. 2) had a value of 0.5 (horizontal bars: 95% CI); α values shifted coherently with changes in A and B values in all four animals, indicating a continuous magnitude-probability trade-off relation. See Fig. 3 for single sessions’ IP values.

The axioms of EUT are necessary and sufficient conditions for choices to maximize Expected Utility (EU). EU = ∑_*i*_ *U*(*m*_*i*_) · *U*(*m*), is defined as the utility *U(m)* of the possible outcome magnitudes (*m*_*i*_) weighted by their respective probabilities of occurrence (*p*_*i*_). This subjectively defined quantity, EU, replaced the objective Expected Value EV = ∑_*i*_ *m*_*i*_ · *p*_*i*_, as a key measure driving decisions (Pascal, 1658).

Compliance with the continuity axiom requires the existence of a probability at which a fixed gamble is choice indifferent against a combination of a higher and a lower gamble with that probability; the axiom implies the possibility of defining a numerical scale of subjective values. Being deterministic rules, the axioms assume perfectly constant preferences over time. In order to account for the stochasticity of choice behavior we interpreted the axioms in a stochastic sense: option A was considered preferred to option B when the proportion of A over B choices was larger than 0.5 (binomial test, *p*<0.05). The standardized logistic regression coefficients were tested for statistical significance through one-sample *t* test. All statistical tests used (binomial test, Spearman rank correlation, one-sample *t* test, likelihood ratio test) were considered significant at *p*<0.05. For multiple comparisons, we applied a false discovery rate (FDR) correction (Benjamini-Hochberg procedure) (Benjamini and Hochberg, 1995). Data analysis was performed using custom scripts in MATLAB (version 8.3.0 (R2014a). Natick, Massachusetts: The MathWorks Inc).

### Task Design

Each of the four animals was trained to express its preferences in > 10,000 trials between pairs of probabilistic reward options, represented as visual cues on a computer monitor. During the experiment, monkeys sat in primate chairs (Crist Instruments) in the laboratory and used arm movements to make choices between two rewarding stimuli presented on a computer monitor. Monkeys A and B moved a joystick (Biotronix workshop, Cambridge) restricted to left/right movements, to control a cursor on a computer monitor vertically positioned 50 cm in front of them; monkeys C and D made arm movements towards a touch-sensitive screen (EloTouch 1522L 15’; Tyco) horizontally mounted at arm-reaching distance. The possible choice outcomes were different amounts of liquid reward (fruit juice), ranging 0.00-0.50ml (Monkeys A and B) or 0.05-0.90 ml (Monkeys C and D). A computer-controlled solenoid valve delivered juice reward from a spout in front of the animal’s mouth. Task event-times were sampled and stored at 1 kHz on a Windows 7 computer running custom MATLAB (The MathWorks) code, using Psychtoolbox 3.

During the initial training period (1-2 months) monkeys learned to select the side of the screen where a single visual stimulus was presented. Initially, the stimulus represented a sure reward, with magnitude varying among up to three levels (between 0.05 ml and 0.5 ml) in pseudo-randomly intermingled trials. Then, to learn the probabilistic information, gambles were introduced in a similar scheme, alternating up to three probability levels (between 0.1 and 0.9). A further training period (2-4 months) included choices between two learned stimuli. Gradually, further magnitude and probability levels and more complex gambles were added to the choice set. Behavioral data acquisition began when preferences were considered stable and clearly differed among the newly introduced gambles.

#### Task design for monkeys A and B

The reward amount (magnitude) was represented though the vertical position of a horizontal white bar within a frame, composed by two thin vertical gray lines. A single option could contain up to three possible outcomes, each with a specific probability. The probability associated with each outcome was cued through the width of the horizontal bar. Each choice option could be either a safe option (i.e. a sure reward or “degenerate gamble”, with probability P=1), presented as a single horizontal bar filling the full width of the frame, or a probabilistic distribution of rewards (i.e. a risky gamble) presented as multiple horizontal bars. The horizontal position of the bars representing non-safe outcomes were randomly shifted horizontally within the frame to avoid that animals only considered a particular portion of the stimulus.

To initiate a trial, the monkey held a joystick in the central position for a variable time interval (1-1.5 s). Two visual cues representing the choice options appeared to the left and right sides of the computer monitor. The monkey indicated the preferred option within 2 s by moving the joystick to the side of one option, at which time the unselected option disappeared. After holding the joystick for at least 1 s, the reward corresponding to the selected option was delivered (Fig. 1a). Visual cues were presented on a blank screen, indicating the amount (magnitude) and probability of receiving a reward (fruit juice) though white horizontal lines: each line’s vertical position indicated a reward amount, while the line width was proportional to the probability of obtaining that reward (Fig. 1b).

#### Task design for monkeys C and D

The reward magnitude was represented by the vertical position of a horizontal black bar within a vertically oriented white rectangle. The probability of a reward was conveyed through a circular stimulus, presented adjacent to the bar stimulus, composed of two sectors distinguished by black-white shading at horizontal and oblique orientation; the amount of horizontal shading indicated the probability of obtaining the cued reward magnitude. On each trial, the animal made a choice between two gambles, one of which was a degenerate gamble (P=1), presented randomly in left-right arrangement on the monitor. For risky gambles, the cued reward magnitude could be obtained with P = cued probability and a fixed small reward (0.05 ml) could be obtained with P = 1 – cued probability.

Each trial started when the background color on the touch screen changed from black to gray. To initiate the trial, the animal was required to place its hand on an immobile, touch-sensitive key. Presentation of the gray background was followed by presentation of an ocular fixation spot (1.3° visual angle). After 500 ms, both choice options appeared in left-right arrangement on the monitor, followed after 750 ms by appearance of two blue rectangles below the choice options at the margin of the monitor, close to the position of the touch-sensitive key. The animal was then required to touch one of the targets within 1,500 ms to indicate its choice. Once the animal’s choice was registered, the unchosen option disappeared and after a delay of 500 ms, the chosen object also disappeared and a liquid reward was given to the acting animal. Reward delivery was followed by a trial-end period of 1,000 ms which ended with extinction of the gray background.

### Logistic regression

To identify the key variables driving choice, we analyzed single trials’ data from each session using the following logistic regression:

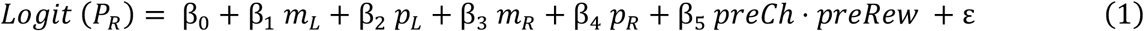

where *P*_*R*_ is the probability of choosing the right-side option; *m* and *p* represent the reward magnitude and probability of options respectively, presented on the left (*L*) or right (*R*) side of the screen; *preCh* represents the previous trial’s choice (−1 for left-side and 1 for right-side choices) while *preRew* corresponds to the reward magnitude obtained in the previous trial; the product *preCh⋅preRew* thus increases for larger rewards obtained when choosing the right-side option in the previous trial; ∊ is the error term. We obtained estimates of the regression coefficients (β_i_) for each session. Coefficients were standardized by multiplying each β with the ratio of the corresponding independent variable’s SD over the SD of the predicted variable. Standardized βs were tested for statistical significance through one-sample t test.

To explicitly investigate variability across sessions, we repeated this analysis on the complete trials dataset (not separated by session), and compared the regression accuracy between the given model (Eq. 1) and a second model which included session number as a random variable. We compared results from the two models (fixed effect and mixed effect) using a likelihood ratio test (p<0.05) and computed the pseudo-R^2^ as a goodness-of-fit measure to compare the accuracy of the two models.

### Axioms of Expected Utility Theory

The axioms of EUT are necessary and sufficient conditions for choices to be described by the maximization of EU: if the axioms are fulfilled, a subjective value corresponding to the EU can be assigned to each choice option, and the option with the highest EU is chosen (von Neumann and Morgenstern, 1944).

Formally,

I. *Completeness*: ∀ A, B either A ≻ B, B ≻ A, or A ~ B
II. *Transitivity*: A ≻ B, B ≻ C ⇒ A ≻ C
III. *Continuity*: ∀ A ≻ B ≻ C, ∃! p ∈ (0, 1) such that pA+(1-p)C ~ B
IV. *Independence*: ∀ A ≻ B ⇒ pA+(1-p)C ≻ pB+(1-p)C; ∀C, ∀p ∈ (0, 1)

Where A, B, C are gambles corresponding to known probability distributions over outcomes, “≻” is the preference relation and “~” represents indifference. The operation pA+(1-p)C corresponds to combining the two gambles A and C with probabilities p and (1-p) respectively, thus representing itself a gamble different from A or C alone.

The continuity of preferences (axiom III) can also be expressed as follows:

III-a. *Monotonicity*: ∀ A ≻ B ⇒ A ≻ αA+(1-α)B ≻ B; ∀ α ∈ (0, 1)
III-b. *Archimedean property*: ∀ A ≻ B ≻ C, ∃! p_1_, p_2_ ∈ (0, 1): p_1_A+(1-p_1_)C ≻ B ≻ p_2_A+(1-p_2_)

Such an alternative expression (III-a, III-b) does not include any equality (i.e. indifference point) and is thus better suited for experimental hypothesis testing compared to III.

Complete (I) and transitive (II) preferences are necessary for univocally and consistently ranking all choice options, representing a “weak ordering” condition. In this case, each possible choice option can be given a specific rank level, so that an option with higher rank will be preferred to one with lower rank. Although these rank levels can be defined as numbers, they have no cardinal meaning: any monotonic transformation of these values would still represent preferences. Such rank levels would give no information about the strength of preferences and could not predict choices between options defined as combinations of gambles.

Conversely, if preferences are also continuous (axiom III), they can have a meaningful numerical utility representation. Thus, if A is preferred to B, the utility of option A (U_A_, a real number) is larger than the utility of B (U_B_) and vice versa if U_A_>U_B_, option A is preferred over option B:

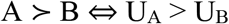

The independence axiom (IV) allows to go one step further, defining how to compute the utility of any gamble G from its attributes (magnitudes *m*_*i*_ and associated probabilities *p*_*i*_):

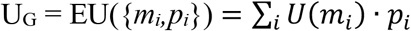

making it possible to predict choices between any possible choice options.

### Expected Utility theorem

Following the four axioms, the EU theorem states that given any two options A and B, A will be preferred to B if and only if the EU of A is larger than the EU of B:

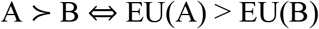

where *EU*(*X*) = ∑_*i*_ *U*(*m*_*i*_) · *p*_*i*_

with X representing a gamble with outcomes *m*_*i*_ and associated probabilities *p*_*i*_ and *U(m)* representing the utility associated with the magnitude *m*. The EU of a gamble thus corresponds to the average utility of a gamble, weighted by the reward probabilities, representing the subjective equivalent to the objective (mathematical) expected value *EV* = ∑_*i*_ *m*_*i*_ · *p*_*i*_.

The EU theorem links preferences to subjective evaluations: if option A is preferred to option B, the EU of option A will be greater than the EU of option B; vice versa, if the EU of A is greater than the EU of B then A will be preferred to B.

### Lexicographic preferences

Lexicographic preferences represent a possible violation of the continuity axiom. Lexicography refers to the way words are ordered based on their component letters: the first letter defines which word comes first in the dictionary, unless words have the same first letter in which case the second letter will define the order, and so on. In choice theory, lexicographic preferences correspond to a decision strategy where the preference for one option is only based on one attribute, while a second attribute is considered only when the first attribute has the same value in both options. In risky choices, the attributes of an option correspond to reward magnitude and probability; in this context, lexicographic preferences imply that the option with the highest magnitude would always be chosen, independent of its probability, unless the two options had the same magnitude, in which case the option with the highest probability would be chosen. Inverting the roles of magnitude and probability would also result in lexicographic choices (Fig. 1g).

Lexicographic preferences, while complying with the completeness and transitivity axioms, represent a violation of the continuity axiom. They imply that reward magnitude and probability are not combined into a subjective value, indicating an underlying choice mechanism (and its neural implementation) incompatible with EUT and with the concept of utility.

### Testing deterministic axioms

To experimentally test an axiom, which is an absolute rule that must hold for any possible gamble, it is necessary to test the largest possible number of different cases. We thus generalized our results by repeating the continuity test using different initial gambles: in one set of tests A, B and C were defined as sure rewards (degenerate gambles) varying over the range 0 to 0.9 ml; in a different set of tests B was defined as a probabilistic two-outcome gamble; in a final set of tests A, B and C were all defined as two-outcome gambles, resulting in the AC option being a three-outcome gamble.

The EUT axioms were originally defined as deterministic rules, which assume that preferences do not change over time. In order to account for the variability in choice behavior (repeated choices between the same pair of options can yield different results) we interpreted the axioms in a stochastic sense: option A was considered preferred to option B when the proportion of A over B choices was larger than 0.5 (P(A≻B)>0.5; binomial test, *p*<0.05).

### Stochastic Transitivity

We tested two stochastic forms of the transitivity axiom, weak stochastic transitivity (WST) and strong stochastic transitivity (SST).

In repeated presentations of the same choices, compliance with WST corresponds to choosing A over B, B over C and A over C in more than 50% of the trials:

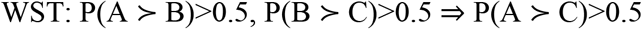

WST thus represents a simple extension of the transitivity rule to stochastic choices.

SST further requires that preference of A over C must be stronger than the other two preferences. Interpreting the proportion of choices for one option as indicating the strength of preference for that option relative to another option, SST can be defined as:

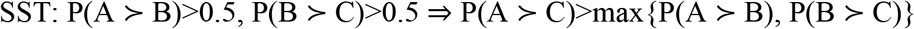

We tested the compliance with WST and SST in choices between different reward amounts. Each test was identified by a triplet of reward magnitudes (degenerate gambles A, B and C), which were then presented in a randomly intermingled sequence of repeated choice pairs (A vs B, B vs C or A vs C). We used a range of possible values for A, B and C rewards: in monkeys A and B, reward A included values between 0.15 ml and 0.50 ml (in 0.05 ml increments), reward B was 0.05, 0.15, 0.20, 0.25, 0.35 or 0.45 ml, reward C was 0, 0.05, 0.15, 0.25 or 0.35 ml. In monkeys C and D, reward A had values between 0.30 ml and 0.90 ml, reward B was between 0.30 ml and 0.90 ml, reward C between 0.10 ml and 0.60 ml (all in 0.10 ml increments). The total number of tested triplets per monkey was: 14 (A), 48 (B), 4 (C), 84 (D).

### Testing the continuity axiom

We implemented a behavioral and statistical test of the continuity axiom as follows:

We defined three starting gambles and verified that monkeys complied with the transitivity axiom; this allowed us to define A, B and C as the most preferred, middle and least preferred gamble respectively.

In each trial, monkeys chose between the middle gamble B and a probabilistic combination (AC) of the most and least preferred gambles: AC = p_A_A+(1-p_A_)C, where p_A_ is the probability of obtaining the most preferred gamble.

We statistically tested compliance with the axiom according to definitions III-a and III-b: we defined a series of AC combinations with specific probabilities (p_A_ between 0 and 1, in 0.1 increments) and measured the proportion of choices for the AC option (P_AC_). We then tested if monkeys preferred the middle gamble to the AC combination for at least one p_A_(P_AC_ < 0.5; binomial test, *p*<0.05, FDR corrected) while also preferring the AC combination to the middle gamble in at least one case (P_AC_ > 0.5), as required by the *Archimedean property* (III-b); compliance with the *Monotonicity* rule (III-a) was ensured by the tested compliance with FSD, and tested in each continuity test by showing increasing preferences for increasing probability of gamble A (rank correlation, *p*<0.05).

After testing for the existence of an indifference point (α), its numerical value was determined by fitting a softmax function to the choice data though non-linear least squares fit. The softmax function was defined as follows:

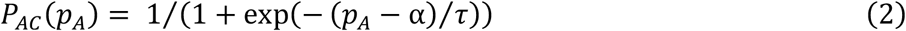

where τ (softmax “temperature” parameter) represents the steepness of the preference function (steeper for lower τ values).

### Data fitting of indifference points

To obtain a set of curves approximating the ICs, for each middle gamble B we fitted the IPs corresponding to varying gamble A magnitudes, using three different functions:

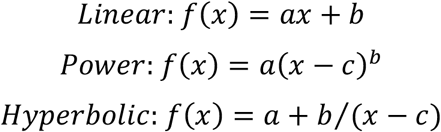

where *x* represents the reward magnitude, *f*(*x*) the IC, i.e. the reward probability as a function of reward magnitude. A non-linear least squares method was used to minimize the error in the probability domain (x-axis in Fig. 4 and Fig. 6b)

### Economic choice models

We modeled the probability of choosing one option using a standard discrete choice model. The probability of choosing gamble A in choices between any two gambles (choice set: {A,B}) was defined through a binary logistic model:

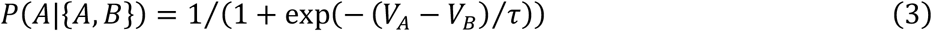

where *V*_*A*_ and *V*_*B*_ represent the subjective values of gamble A and B respectively, τ the temperature parameter. The gamble value was defined following EUT as *V* = *EU* = ∑_*i*_ *U*(*m*_*i*_) · *p*_*i*_, with utility *U*(*m*) being a parametric function of reward magnitude *m*.

A maximum likelihood estimation (MLE) procedure was used to estimate the free parameters (τ and utility-function parameters) to best approximate the measured proportions of choices.

The MLE procedure involved computing and maximizing the log-likelihood (*LL*) in the parameters space.

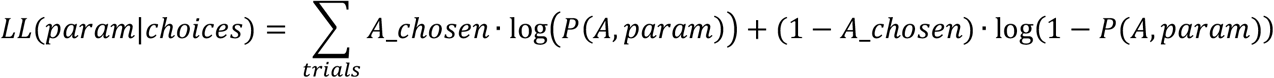

where the sum is defined across all trials in one session, *A*_*chosen*takes the value of 1 if gamble A was chosen in one trial, zero otherwise, and *P*(*A*, *param*) is the discrete choice model defined above with parameters *param*. We minimized the negative LL using the *fminsearch* Matlab function.

We defined three possible utility functions:

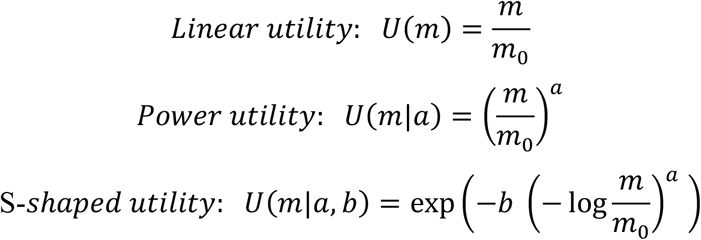

with *m*_0_ = 0.5 *ml*, representing the maximum reward magnitude, thus normalizing all utility functions between 0 and 1.

A linear utility function could only explain choices based on the expected value of the options: it would perform as the best model only if monkeys were choosing by comparing the objective, mathematical expected value of the options. A power utility function would instead be able to describe choices with a specific risk preference: either risk seeking or risk aversion. Finally, an s-shaped utility function could accommodate a more complex pattern of risk attitudes, with the possibility of both risk seeking and risk aversion for different reward magnitudes. As the s-shaped function we used the two-parameter *Prelec* function, which is typically used as a probability weighting function, but can also represent a plausible shape for the utility function.

Using the same binary logistic model with a different definition of the gamble value allowed us to test models from different economic choice theories. In prospect theory (PW model) the gamble value is defined as *V* = *U*(*m*) · *w*(*p*); we used the two-parameter *Prelec* function as the probability weighting function *w*(*p*). In the additive model we defined a gamble’s value as *V* = *ω*_*m*_ · *U*(*m*) + *ω*_*p*_ · *p*, with *ω*_*m*_ and *ω*_*p*_ as additional free parameters acting as “weights” for the magnitude and probability terms. According to the mean-variance approach, the value definition does not rely on the utility concept: *V* = *EV* + β · *Risk*, where *EV* and *Risk* are the first two moments of the gamble’s probability distribution, and β is a free parameter. We computed *Risk* as the expected value of the squared deviation from the mean: *Risk* = ∑_*i*_ *p*_*i*_ · (*m*_*i*_ − *EV*)^2^.

In order to construct the full indifference map predicted by a model, for each IC we numerically computed the indifference points corresponding to finely spaced magnitude levels: for a selected model (using the average recovered parameters across all sessions), the subjective value of the B gamble was computed (V_B_); after increasing the magnitude by 0.001 ml (starting from the B gamble magnitude), the subjective value was then computed for a series of probabilities (step 0.001), and the probability corresponding to the value closest to V_B_ was identified as the indifference point. This procedure, repeated for all B gambles, allowed us to obtain a distance measures between all the modeled and measured IPs, in the probability domain, which was used as one of the quantities for model comparison.

To compare the six tested models (EUT with three possible utility functions, PW, additive model and mean-variance) we defined four accuracy metrics: 1) the square root of the mean squared error, representing the average distance between modeled and measured IPs, in probability units; 2) the Bayesian information criterion (BIC) and 3) the Akaike information criterion, both introducing a penalty term when increasing the number of model parameters; the variance in the differences of modeled vs measured preferences, i.e. the proportion of AC vs B choices across all continuity tests. For each of these four measures, a lower value represented a better model compared to a higher value.

## RESULTS

### Rationale

In EUT, decisions are modeled “as if” subjects had an internal utility representation, making no assumptions about the brain processes underlying choice (Camerer, 2008). To investigate the possibility of EUT describing the actual neuronal mechanisms of choice, our approach is to 1) verify that subjects follow the model’s assumptions, 2) infer the subjective utility measure defined in EUT, which is not directly measurable and 3) identify the neuronal substrates coding such quantity, if they exist. To fulfill the first point, we need to verify compliance with the assumptions of EUT (i.e. the axioms). If the assumptions are satisfied, the utility measure can be elicited following econometric methods. These crucial steps identify the subjective quantities, as opposed to the objective, physical ones, that can be used to describe preferences. The third point, which represents the ultimate goal of our research, involves the identification of utility-coding neuronal substrates by correlating the neuronal activity with the utility measure rigorously defined in the previous points.

Here, we focused on the initial step: testing the basic assumptions of EUT in order to infer the existence of a utility measure. We based our experimental tests on the third EUT axiom, the continuity axiom, which is crucial for defining a numerical utility measure. The fourth axiom (independence) is needed to mathematically define the subjective value as an option’s EU. Violations of the independence axiom have been reported in human and animal experiments, highlighting the limits of EUT (Allais, 1953; Kahneman and Tversky, 1979; Battalio et al., 1985; Conlisk, 1989). Nevertheless, the continuity axiom remained a necessary condition in all major generalized EU theories developed since the 1940s, which share the axiom’s main implication, i.e. the definition of a scale of numerical subjective values (Harless and Camerer, 1994; Starmer, 2000; Jehle and Reny, 2001).

We carried out our study on macaque monkeys, a close evolutionary neighbor to humans, who are able to perform skilled problem solving after relatively short training periods. For each animal, it was possible to elicit hundreds of choices per day, for several months. This allowed us to systematically investigate a broad range of choice problems, with a fine resolution in the reward magnitude and probability domains, which is difficult to achieve in typical economic studies on humans. In particular, the idea of testing an axiom only makes sense if we are able to test compliance with it in an extensive and comprehensive set of choice situations. Indeed, this is what drove our definition of the choice sets in the current experiment: extending the axiom test to gradually more complex choice cases, with the highest attainable resolutions. As a downside, due to the high experimental costs and long durations of primate research, we achieved a lower number of participants compared to typical human studies. We believe that this limitation was counterbalanced by the robustness of the results that were obtained in each animal, thanks to the elevated number of repetitions and choice problems tested. The currently developed systematic choice task should stimulate future neuronal investigations of decision variables in primates using well established formal economic theory.

### Design

To test the continuity axiom of EUT in non-human primates, we trained four monkeys to perform a binary choice task. In each trial, the animal chose between two options, presented simultaneously on a computer monitor (Fig. 1a), offering liquid rewards varying in amount and probability (Fig. 1b). The continuity axiom states that given any three ranked gambles (A, B and C, ranging widely) a decision maker should be indifferent between the middle gamble (B) and a probabilistic combination of the two other gambles (AC). Formally,

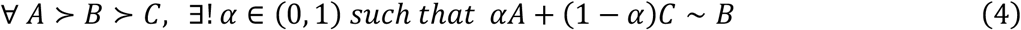

where “≻” defines a preference relation and “~” indifference; *α* is the specific probability associated to gamble A for which indifference occurs. Note that the axiom should be satisfied for any arbitrary set of gambles A, B and C.

The axiom was originally defined as a “plausible continuity assumption” (von Neumann and Morgenstern, 1944). Thought experiments intuitively clarified how the continuity axiom could be violated (Georgescu-Roegen, 1954; Levin, 2006; Chateauneuf et al., 2008), for example when options had infinitely different values. Nevertheless, the axiom was considered a reasonable condition and not experimentally tested.

To experimentally test the axiom, we first defined three gambles (Fig. 1c) for which the monkey had well defined preferences (A≻B and B≻C in the majority of trials; binomial test, *p*<0.05). We then combined the most and least preferred gambles (A and C respectively) with probability pA (varying between 0.1 and 0.9 in 0.1 steps), obtaining the family of gambles AC(p_A_) (Fig. 1d). Finally, we presented choices between B and one of the AC combinations and tested for the existence of indifference between B and a probabilistic combination of gambles A and C, with probability *p*_*A*_ = *α* such that *B* ~ *αA* + (1 − *α*)*C* (Fig. 1e). Compliance with the continuity axiom would thus be demonstrated by the existence of a unique α between 0 and 1 (Fig. 1f), while violations would occur if α were not identifiable or when multiple α existed (Fig. 1g).

Therefore, we varied the behavioral test with the A, B and C gambles in several ways: (*1*) we varied the safe reward amounts of the degenerate gambles A, B and C between tests (see paragraph *Compliance with the continuity axiom*); (*2*) we used a risky B gamble but kept varying the safe reward amounts of the degenerate gambles A and C between tests (see paragraph *Indifference curves in the magnitude-probability space*); (*3*) we used only risky A, B and C gambles and varied A and C between tests (see paragraph *Continuity axiom test in the Marschak-Machina triangle*). The first, more basic manipulation (*1*) was tested in four monkeys, while the further two, more specific variations (*2* and *3*) were tested in only two of the animals (monkeys A and B).

We used pseudo-random repetitions of all presented choice pairs in order to account for the stochasticity of choice behavior and as a basis for future recordings of neuronal activity. Although the EUT axioms were originally defined as deterministic rules (von Neumann and Morgenstern, 1944), we extended their definition to the stochastic domain (Hey and Orme, 1994; Loomes, 2005) because the statistical analysis of neuronal responses requires the use of multiple trials. Therefore, we made our design compatible with basic assumptions of stochastic choice theories (Luce, 1959; McFadden and Richter, 1990; McFadden, 2005). We quantified the animal’s choice from the probability of choosing one option over its alternative, rather than employing traditional single-shot economic tests.

### Basic choice behavior

We investigated the consistency of choice behavior to make sure that the four tested monkeys understood the reward-cue associations and were able to express their preferences.

To assess the contribution of magnitude and probability information to decisions, we performed a logistic regression on single trials’ choice data, using the chosen side as the dependent variable and the options’ probabilities and magnitudes as independent variables. An additional regressor controlled for the effect of past trials: the product of the previous trial’s chosen side and obtained reward (Methods section: *Logistic regression*, Eq. 1). Standardized regression coefficients (β) corresponding to reward magnitude and reward probability were significantly different from zero (one-sample *t* test, *p*<0.05, FDR corrected) in all four animals (Fig. 1h), indicating that both variables were choice-driving factors. Compared to such coefficients (average absolute value across animals: 0.54±0.18 SD), the past trials’ β was much smaller (average absolute value: 0.032±0.025 SD) and not consistently significant across animals, confirming that choices were mainly driven by the two cued attributes. A significant intercept implied a side bias for monkeys A, B and C (one-sample *t* test, FDR corrected; per animal: *p*=1.7·10^−10^ (A), *p*=5.1·10^−3^ (B), *p*=7.3·10^−5^ (C), *p*=4.1·10^−1^ (D)). The side bias was accounted for by presenting each option on both sides of the screen, the same number of times.

As a direct test of consistent choice behavior, we verified compliance with first order stochastic dominance (FSD). FSD is the probabilistic analogue of “more is better”, and represents a basic requirement of EUT and of the continuity axiom in particular (Methods section: *Axioms of Expected Utility Theory*): a gamble should be preferred if it contains outcomes at least as good as another gamble, with at least one strictly better outcome. FSD implies that an option with a more probable outcome should be preferred to one with a less probable outcome of the same reward amount. The higher probability gamble stochastically dominates the lower probability gamble and should thus be preferred. Due to choice stochasticity, a number of dominated choices are naturally expected, but to comply with FSD their proportion must be significantly below 0.5. We tested FSD in choices between a gamble and a safe option as well as between two gambles, using reward magnitudes (fixed for each presented pair of options) between 0.1 and 0.9 ml and reward probabilities ranging from 0.05 to 0.97 (step 0.02, monkeys A and B) and from 0.25 to 0.75 (step 0.125, monkeys C and D). We found that all four animals complied with FSD by preferring the dominant option in more than 50% of trials across all FSD tests (Fig. 1i; binomial 95% CI above 0.5). We inferred from this behavioral compliance with FSD that the animals attributed higher reward value to higher reward probability, as prerequisite for testing the integration of reward probability and magnitude with the continuity axiom.

A further prerequisite for testing the continuity axiom is compliance with the completeness and transitivity axioms. Completeness ensures that subjects have well-defined preferences for any presented pair of options. In line with general notions of discrete choice models (McFadden, 2001), in every trial our choice set had a finite number of offered alternatives (two) with mutually exclusive (only one option could be selected) and collectively exhaustive options (the set included all possible options). Animals were thus induced to express complete preferences. Still, they could choose not to select any option, avoiding expressing a preference, which would violate the completeness axiom. This was not consistently observed, except rarely for low-valued options pairs (which were excluded from subsequent testing). Thus, we tested the animals’ choices while they complied with the completeness axiom.

The transitivity axiom ensures that all gambles can be univocally ranked. In line with stochastic choice theory, we tested two stochastic forms of transitivity, weak (WST) and strong (SST) stochastic transitivity (Methods section: *Stochastic transitivity*), using combinations of the A, B and C magnitudes ranging from 0 to 0.5 ml (step: 0.05 ml) for monkeys A and B, and from 0 to 0.9 ml for monkeys C and D (step: 0.1 ml). In choices from all tested triplets, the four animals complied with both WST and SST (Fig. 1j). Individual transitivity tests revealed compliance with WST in all 141 tested magnitude combinations and compliance with SST in 125 (89%) tested triplets (average number of trials per test, per animal: 21 (A), 96 (B), 36 (C), 105 (D)). This compliance with the transitivity axiom indicated that the animals made consistent choices and thus ranked the tested gambles unequivocally.

### Compliance with the continuity axiom

Following the formal definition of the continuity axiom (Eq. 4), we assessed the existence of a unique IP in choices between a fixed gamble and a probabilistic combination of the other two gambles. We defined three degenerate gambles A, B and C with three different reward amounts; in each trial, the animal chose between the middle gamble (B) and the probabilistic combination of the most and least preferred gambles (A and C, respectively). Thus, we tried to obtain a p_A_ at which choice indifference occurs: α=p_A_ such that B ~ p_A_(A) + (1-p_A_)C.

All four animals preferred the middle gamble to at least one of the AC combinations, while also preferring at least one of the AC combinations to the middle gamble (Fig. 2): for different pA values, the proportion of choices for the AC combination was significantly below or above 0.5 (binomial test, *p*<0.05) (Fig. 2a,c), following an increase in preference with increasing pA (rank correlation, *p*<0.05). Such a switch of revealed preference depending on probability pA indicated the existence of a unique IP and thus compliance with the continuity axiom (Methods section: *Testing the continuity axiom*).

We defined the A, B and C gambles as degenerate gambles of varying reward magnitudes. All tested triplets showed a pattern of AC preferences compatible with the continuity axiom: the existence of both preferred and non-preferred AC combinations, together with gradually increasing preferences of the AC option, implied the presence of an IP. Monkeys complied with the continuity axiom when defining the C gamble as 0 ml (monkeys A and B, Fig. 2a,b) as well as when using a non-zero C gamble (monkeys C and D, Fig. 2c) in a different task (Methods section: *Task Design*) thus confirming the robustness of our results.

Importantly, different triplets of gambles produced IPs varying in a meaningful and consistent manner: increasing only the reward magnitude of the middle gamble (B) (e.g. from 0.25 ml to 0.35 ml; Fig. 2a,b) produced larger α values; decreasing the magnitude of the most preferred gamble (A) (e.g. from 0.50 ml to 0.40 ml (Fig. 2b)) resulted in higher α values (non-overlapping 95% CI, *p*<0.05). Such a pattern reflected the notion, central to the continuity axiom, of α being a measure of the subjective value of the middle gamble: the more B was considered close to A in value, the higher its α; the closer B was considered to C, the lower its corresponding α. While shifting consistently for different initial gambles, the α values were also different across animals, denoting the subjective quality of the measured IPs. In conclusion, testing the continuity axiom showed a coherent pattern of IPs, highlighting the joined contribution of reward magnitude and probability to the definition of subjective values.

Lexicographic preferences represent a possible continuity axiom violation (Fig. 1g). Lexicography refers to the way words are ordered based on their component letters. In analogy, lexicographic preferences in risky decision making correspond to choices based on one component at a time (either reward magnitude or probability). They represent a specific choice heuristic in which the gamble components are considered separately and are not combined into a single quantity. This corresponds to a choice mechanism incompatible with the definition of numerical subjective values: lexicographic choices cannot be described by assigning a numerical value to each gamble, as in EUT (Methods section: *Lexicographic preferences*). By showing the existence of a coherent set of IPs, our data demonstrated that preferences were not lexicographic, implying that animals considered and combined magnitude and probability information.

Although compliance with the axiom was robust when considering the average IPs, there appeared to be a clear variability of IPs across sessions. When looking at the variation of the IPs over time (extended data Fig. 3a), we found that they showed a combination of a random variation and linear trend periods. This was evident in monkey B’s choice behavior, where different sessions produced clearly different IP values for the same continuity test (extended data Fig. 3b). This resulted from values gradually varying by session, reflecting gradual changes in the risk attitude over long periods of time (days/months), possibly due to factors external to the experiment-defined variables (including learning of the reward-stimuli association and adaptation to uncontrolled variables). Importantly, the observed variability did not affect the ability to robustly identify the IP in a single session (Fig. 3b), a necessary requirement for the possibility of correlating such behavioral measures with the activity of single neurons or neuronal populations.

**Figure 3.**
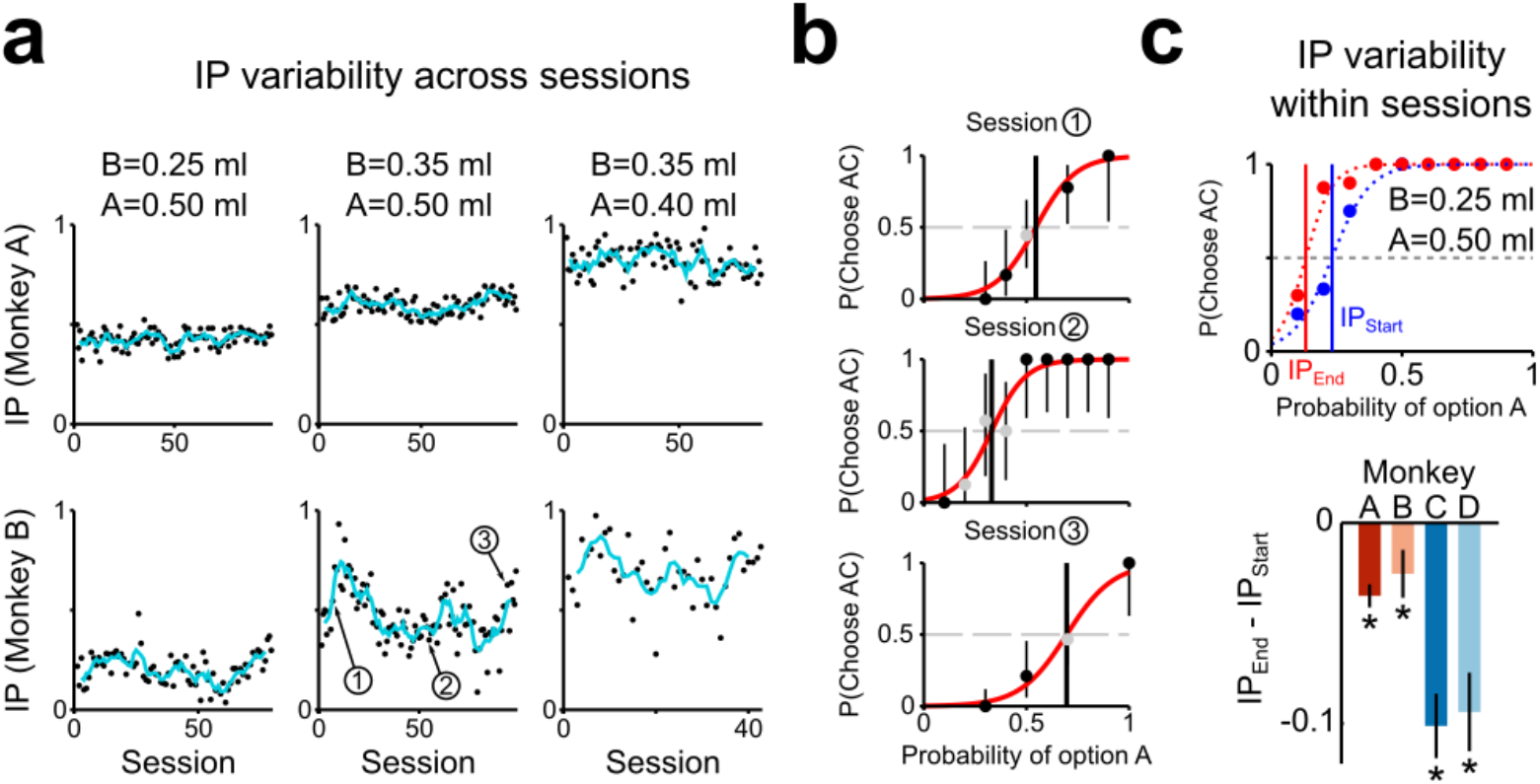
Variability of IP across and within sessions. **a**) Evolution of the IP as a function of session number (blue curve: 5-session moving average), for the three continuity axiom tests presented in Fig. 2a,b (Monkeys A, B). **b**) Choice proportion for the AC gamble in three example sessions (as indicated in panel a). Vertical bars: binomial 95% CI, indicating a significant preference (black dot) when not crossing the horizontal dashed line. **c**) Top: preferences in the first half (blue) and in the second half (red) of a session (one example session, Monkey B); bottom: average IP change across all sessions (bars: SE). Asterisk: IP difference significantly different from zero (t test, *p<0.05*).

By including the session number as a random variable to the previously defined logistic regression we compared the performance of a mixed effect model (including both fixed and random effects) with that of the basic fixed effect one (Eq. 1). Adding the random effect resulted in a better description of choices (likelihood ratio test, *p<0.05*), confirming that random variations of choice-driving parameters were observed across sessions. Nevertheless, the increase in the goodness of fit compared to the model not containing the random effects was only marginal (difference in pseudo-R^2^ between the mixed effect model and the fixed effect model, per monkey: 0.01 (A), 0.02 (B), 0.02 (C), 0.01 (D)) with no evident differences in the regression coefficients.

When looking at the variability within single sessions, by measuring the IP difference between the last half and the first half of each session, we found that the IPs significantly decreased over one session’s duration (paired t-test, *p<0.05*) in all four monkeys (extended Fig. 3c). This result suggested the contribution of the total fluid intake as one of the factors affecting IPs, although its effect-size (mean IP difference, per monkey: −0.03 (A), −0.02 (B), −0.10 (C), −0.10 (D)) was smaller than the smallest intervals of probability values used (0.10 for monkeys A, B; 0.125 for monkeys C, D).

Overall, these results support the core ideas arising from the continuity axiom: subjective values, which define preferences, are quantities (numbers) that depend on reward magnitudes and are modulated by reward probabilities. In other words, probabilities modify the subjective reward values in a graded and continuous way; a variation in reward magnitude can be compensated by a change in reward probability and vice-versa, establishing a continuous trade-off relation between magnitudes and probabilities.

### Indifference curves in the magnitude-probability space

To confirm the existence of a continuous trade-off relation between reward magnitudes and probabilities, as implied by the continuity axiom, we represented the animals’ IPs (measured through a softmax fit, Eq. 2) in a two-dimensional diagram with reward magnitude and probability (MP) as variables. Such MP space was used to represent the continuity axiom tests, carried out in monkeys A and B, in which the B gamble is either a degenerate gamble or a true (non-degenerate) gamble, with degenerate A and C gambles (C = 0 ml). Each gamble used in a continuity test (B and AC combinations) corresponded to a single point in the MP space (Fig. 4a). Compliance with the continuity axiom was manifested as choice indifference between the B gamble and an AC combination, identifying a single point in the MP space (B~AC in Fig. 4a).

**Figure 4.**
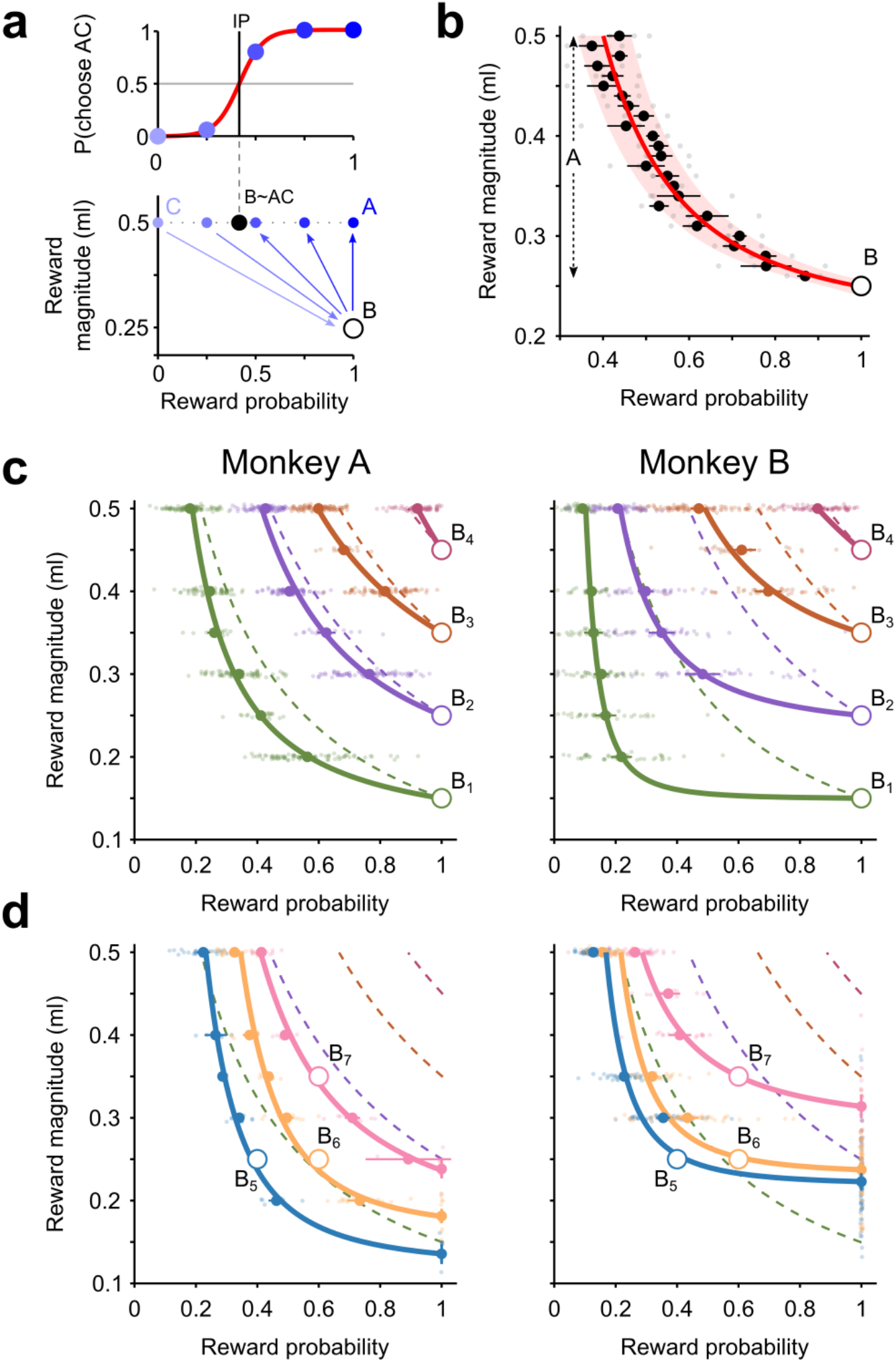
Indifference curves in the magnitude/probability (MP) space. **a**) **Representation of the continuity axiom test in the MP space**. The gambles used for testing the axiom can be mapped into the magnitude-probability diagram. Preference in choices between B (circle) and combinations of A and C (graded blue dots) is represented by an arrow pointing in the direction of the preferred option (bottom), consistently with the proportion of choices for the AC option (top). Each continuity axiom test resulted in an indifference point (vertical line, top), represented as a black dot in the MP space (bottom). **b**) **Indifference curve (IC)**. IPs (gray dot: single session; black dot: average across sessions; bar: SE) obtained using different A values (step 0.01 ml) shifted continuously, producing an IC in the MP space (area: 95% confidence interval). Curve: best fitting power function. Data from monkey A (5 sessions, 1781 trials). **c,d) Indifference map**. ICs for different B values (colored thick curves), described the gradual variation of the average IPs (colored dots, with SE bars) for each B. ICs were modeled as power functions (see Table 4-1 for comparison with hyperbolic and linear fits). Small dots represent IPs measured in single sessions. Both sure rewards (**c**) and probabilistic gambles (**d**) as B options, produced coherent indifference maps, with smooth and non-overlapping ICs. Dashed curves represent points with the same objective expected value, highlighting the subjective quality of ICs.

**Table 4-1.** Comparison of indifference curves’ fitting models. Mean squared error (MSE) resulting from fitting IPs to three different models. Bold face indicates the lowest MSE value for each animal.

| *Model* | *Linear* | *Hyperbolic* | *Power* |
| --- | --- | --- | --- |
| Monkey |  |  |  |
| A | $1.1 \cdot 10^{-2}$ | $9.9 \cdot 10^{-5}$ | **$9.6 \cdot 10^{-5}$** |
| B | $3.4 \cdot 10^{-2}$ | $2.2 \cdot 10^{-4}$ | **$1.7 \cdot 10^{-4}$** |

To test compliance with the continuity axiom for an extended set of degenerate gambles, we held the B gamble fixed but varied the magnitude of the A gamble. This test yielded a set of IPs that lined up as an indifference curve (IC). Importantly, there were no discontinuities (‘jumps’) in the IC while varying the A magnitude in 0.01 ml steps. This was confirmed through a leave-one-out regression, by measuring the percentage of left-out data points falling within the CI identified by all other points: 96% of average IPs (24 out of the 25) and 94% of single session’s IPs (72 out of 77) fell within the 95% CI of the nonlinear IC regression (power function, see below). Such very gradual change in the IP fulfilled a fundamental requirement of the continuity axiom: as the magnitude of the A gamble increased, IPs gradually decreased without any apparent discontinuity (Fig. 4b).

Repeating the IC elicitation procedure for different degenerate B gambles yielded a set of ICs, i.e. an indifference map, which captured the full pattern of relations between reward magnitudes and probabilities. To measure each animal’s indifference map we performed 14 continuity axiom tests, by systematically varying the magnitudes of gambles A and B between 0.15 and 0.50 ml in 0.05 ml steps. For each middle gamble (B_1_ to B_4_) we varied the value of gamble A, obtaining a set of IPs in each session (average sessions per continuity test, per animal: 64 (A), 48 (B)), thus confirming the compliance with the continuity axiom for a large set of A and B magnitudes. We modeled the resulting IC through a power function, which we identified as the best fitting function compared to linear and hyperbolic ones (Fig. 4-1). The fitted IC followed the gradual shift in IPs observed when varying the reward magnitude of gamble A. The indifference map, obtained by including ICs corresponding to all tested B gambles, captured the full pattern of relations between IPs, highlighting their smooth and continuous transitions (Fig. 4c).

As the EUT axioms should apply to any arbitrary set of gambles, we further tested compliance by using a set of truly risky B gambles (B_5_ to B_7_). These 14 experimental tests involved choices between pairs of probabilistic gambles with no option of getting a sure reward, making it a more general and more complex choice situation (average sessions per continuity test, per animal: 12 (A), 40 (B)). Nevertheless, IPs were still consistently observed, and the resulting ICs had qualitatively similar shapes to the ones involving a degenerate gamble (Fig. 4d).

Altogether, these results confirm compliance with the continuity axiom in a broad class of choice situations and highlight the existence of an orderly trade-off relation between reward magnitudes and probabilities: even a small decrease in reward magnitude was compensated in revealed preference by an increase in reward probability and vice-versa.

### Economic modeling of indifference curves

We investigated whether our results were compatible with theoretical economic models of choice in the framework of EUT, particularly in relation to the existence of a utility function able to represent choices in agreement with the EU theorem. According to EUT, a gamble’s value stems from the product of the reward’s utility and its associated probability; this assumes the existence of a utility function over magnitude values, which fully defines the shape of the whole indifference map, uniquely identifying the subjective magnitude-probability trade-off relation (Fig. 5). Note that assuming a linear utility function results in choices depending only on the objective quantities: the EU model incorporates the objective EV model, which represents the objectively optimal preference pattern (Fig. 5a), as a special case.

**Figure 5.**
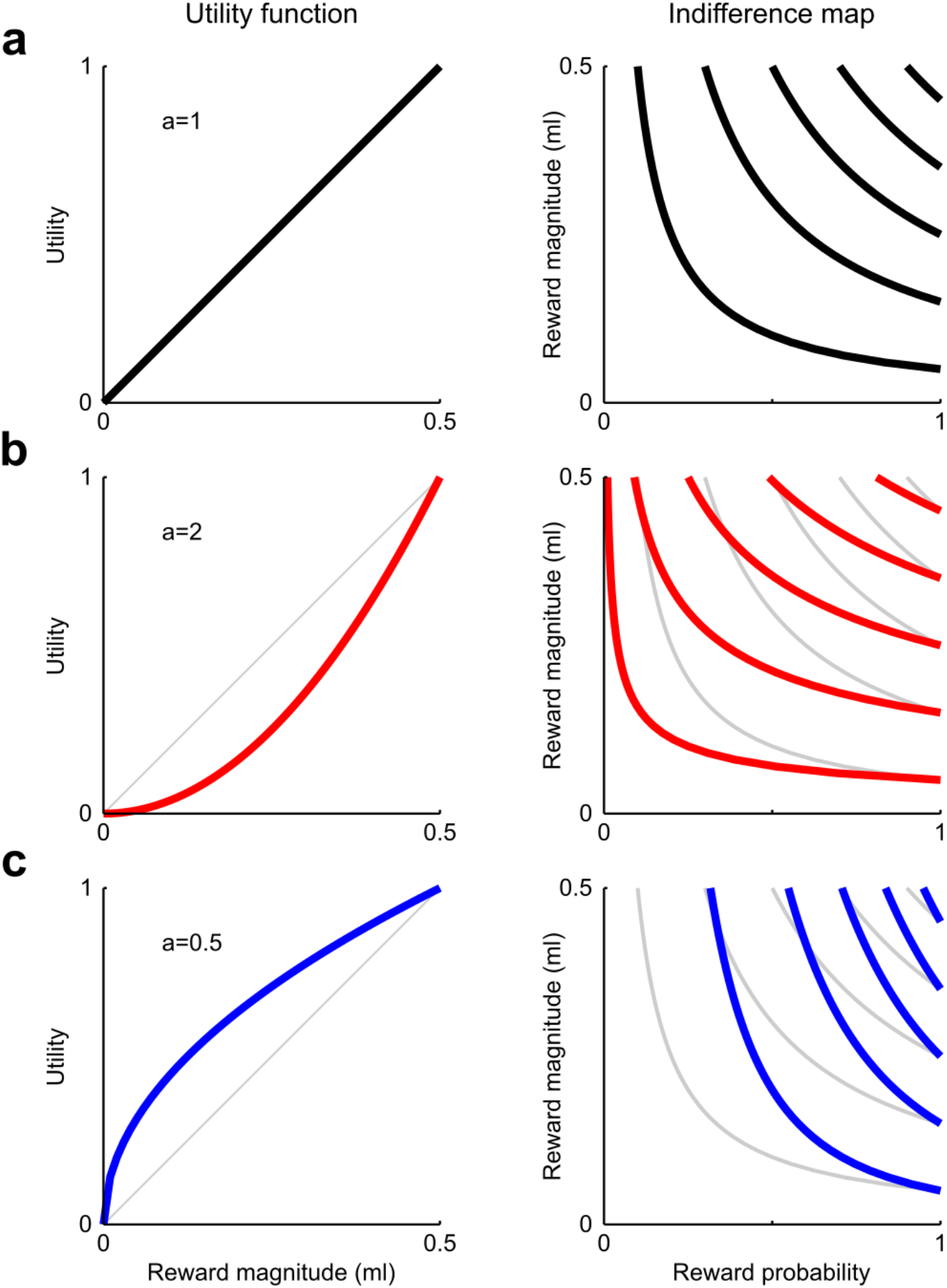
Theoretical relation between utility function and indifference map. Sample indifference maps obtained from different utility functions, with *a* representing the single parameter of a power function (*U*(*m*) = (*m*/*m*_0_)^*a*^, with *m*_0_ = 0.5 *ml*). The indifference curve for a degenerate gamble B was analytically obtained as a function of magnitude values, from the equation *EU*_*B*_ = *U*(*m*) · *p*. By solving the equation for p, the indifference curve equation can be obtained: *p*(*m*) = (*m*_*B*_/*m*)^*a*^, where *m*_*B*_ is the magnitude of a degenerate gamble B. The three utility shapes are directly related to different risk attitudes: risk neutrality for linear utility (a), risk seeking for convex utility (b), risk aversion for concave utility (c). In all plots, risk neutrality (i.e. choices based on the objective EV of gambles) is represented by grey curves. The indifference map globally warps according to the risk attitude.

We estimated the utility function using single-trial choices from each session, through a maximum likelihood estimation (MLE) method. We defined a discrete choice model in standard fashion (McFadden, 2001), with the probability of choosing one option described by a logistic function (Eq. 3), dependent on the difference in EU between the two options. Each gamble’s EU was computed as the utility of the reward multiplied by its probability (Methods section: *Economic choice models*).

We compared MLE results from three utility functions: linear, power and s-shaped. The power utility function captured the monkeys’ choice behavior better than the linear one (difference in Bayesian Information Criterion, BIC: 51.7±39.1 SD, *p*=2.9·10^−18^, Monkey A; 54.9±29.5 SD, *p*=1.7·10^−27^, Monkey B; one-sample *t* test), while the s-shaped utility function outperformed the power-shaped one (BIC difference: 12.2±12.3 SD, *p*=2.1·10^−13^, Monkey A; 8.7±12.2 SD, *p*=1.4·10^−8^, Monkey B). The two recovered parameters for the s-shaped utility functions (Fig. 6a, histograms) were both significantly different from one (*p<*10^−15^ in both monkeys; one-sample *t* test), confirming that utility functions were non-linear and had a significant inflection point (i.e. a change in curvature), resulting in an s-shaped curve.

**Figure 6.**
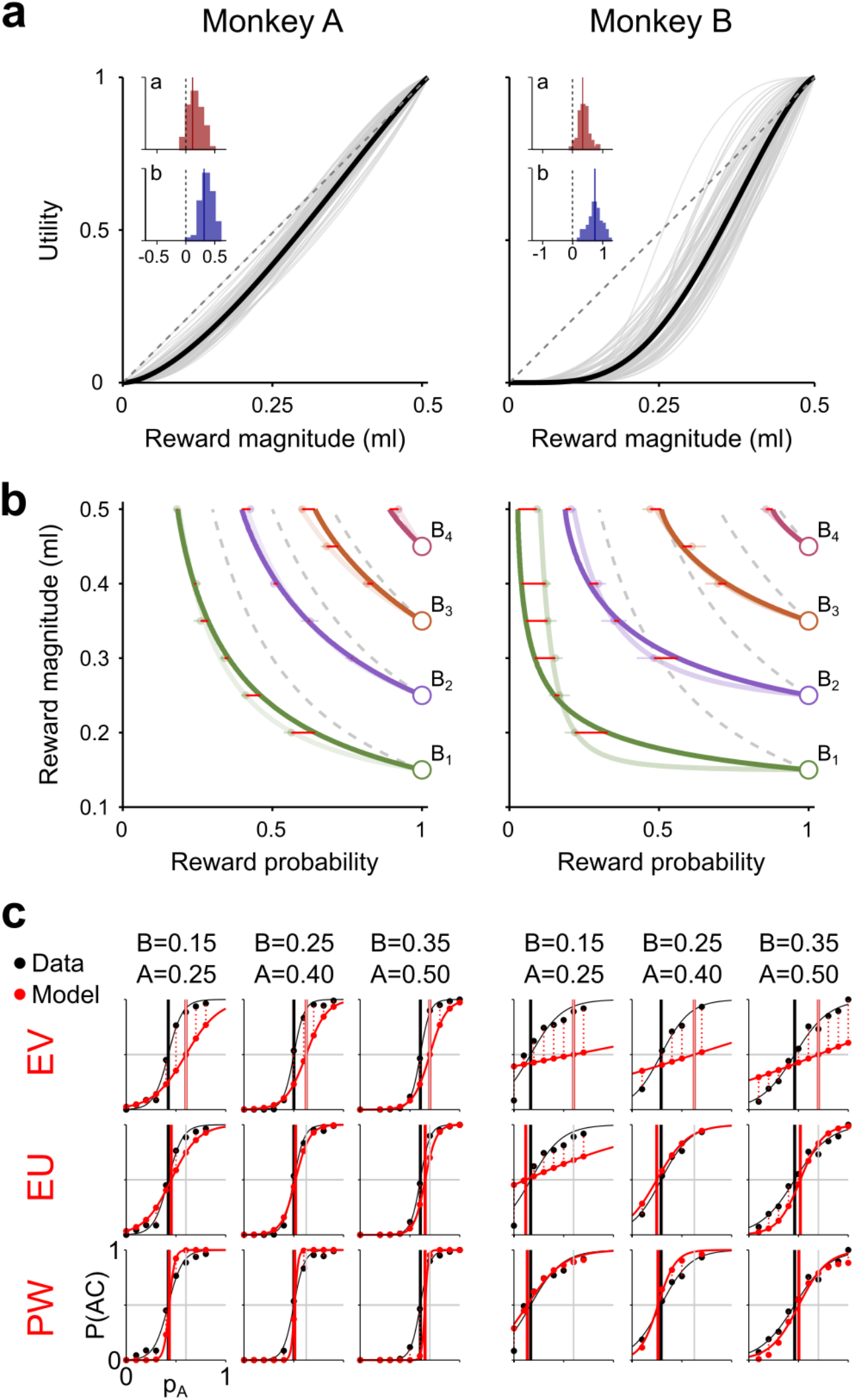
Indifference curves are compatible with economic models. **a) Utility functions.** Single sessions’ utility functions (gray) and averages (black) estimated through MLE using single-trial choice data. The two estimated parameters (*a* and *b* in inset, histograms of log-values) were both significantly positive, indicating s-shaped utility functions. **b) EUT-predicted indifference curves (ICs)**. Indifference map reconstructed using the estimated utility functions. The light-colored curves represent the measured indifference map (Fig. 4c); the red horizontal lines identify the distance between measured and modeled indifference points (IPs). Dashed gray curves define points with equal EV, corresponding to a linear utility model. See Fig. 5 for the theoretical relation between the utility function and the ICs. **c) Comparison of modeled and revealed preferences**. Percentage of choices for the AC gamble (P(AC)) measured (black) and modeled using three models (red), for three example A-B-C triplets (top, A and B in ml, C=0 ml). The EV model could only predict IPs equal to the EVs (grey vertical lines), with a larger error in the prediction of the P(AC) (vertical dotted red lines) compared the EU model. The PW model, which included a subjective probability weighting, was better at capturing the revealed preferences only in specific cases (e.g. B=0.15, A=0.25 in Monkey B).

We used the recovered s-shaped utility functions (Fig. 6a) to construct the corresponding indifference map: for each gamble B we computed its EU and obtained an IC as the set of points with equal EU in the MP space. It was thus possible to define a whole indifference map using a single utility function. Such a map, modeled from the MLE-estimated utility function, closely matched the behavioral IPs and the previously fitted ICs (Fig. 6b), which had been measured for each B gamble independently and had no link to the economic theory. The average distance between the modeled IPs and the behavioral IPs (red lines in Fig. 6b) was smaller for the EU model than for the objective-EV model (dashed curves in Fig. 6b) (square root of the mean squared error: 0.028 (EU model), 0.108 (EV model) for Monkey A; 0.052 (EU model), 0.274 (EV model) for Monkey B). Thus, the EU model was better at capturing the shape of the indifference map compared to the objective EV model by 3.9 times in Monkey A and 5.3 times in Monkey B. We quantified the ability of the model to describe the actual preferences (proportion of choices for the AC option), from which the IPs were calculated, using the variance in the deviation between predicted and measured proportion of choices (vertical dotted lines in Fig. 6c). A lower average variance for the EU model indicated that it was better at describing preferences compared to the EV model (*Var*, Table 6-1). This was also confirmed using a standard model comparison analysis (*BIC* and *AIC* scores, Table 6-1). These results indicate that the non-linearity in the utility function was able to capture the subjective quality of the IPs (Fig. 6b) and of the revealed preferences (Fig. 6c).

**Table 6-1.** Numeric comparison of economic models. Each row of values is a comparison across models using one regression model accuracy metric (averaged across all tests and sessions), with bold face indicating the best fitting model according to that metric (gray font for the worst fitting one). EV corresponds to the EU model assuming a linear utility function. In the PW model the gamble value was computed as *V=U(m)· w(p)* (valid, as defined in Prospect Theory, for all gambles with one non-zero outcome), with *w(p)* being the probability weighting function (2-parameter *Prelec* function). In the additive model, *V=wm·U(m)+wp·p*. The square root of the MSE represents the average distance between model and IP in probability units. *Var* represents the variance in the differences of modeled vs measured preferences (the proportion of AC vs B choices across all continuity tests).

|  | *Model* | *EV*<br>*U: linear*<br>*w: linear* | *EU*<br>*U: power*<br>*w: linear* | *EU*<br>*U: Prelec*<br>*w: linear* | *PW*<br>*U: power*<br>*w: Prelec* | *Mean-*<br>*variance* | *Additive 1*<br>*U: Prelec*<br>*w: linear* | *Additive 2*<br>*U: power*<br>*w: Prelec* |
| --- | --- | --- | --- | --- | --- | --- | --- | --- |
|  | *Metric* |  |  |  |  |  |  |  |
| Monkey A | $\sqrt{MSE}$ | 0.108 | 0.046 | 0.028 | **0.024** | 0.066 | 0.091 | 0.126 |
|  | *BIC* | 348 | 296 | **293** | 294 | 308 | 347 | 320 |
|  | *AIC* | 344 | 288 | 281 | **275** | 300 | 331 | 301 |
|  | *Var* | 0.154 | 0.122 | 0.117 | **0.113** | 0.129 | 0.142 | 0.125 |
| Monkey B | $\sqrt{MSE}$ | 0.274 | 0.093 | **0.052** | 0.062 | 0.193 | 0.212 | 0.084 |
|  | *BIC* | 379 | 324 | 316 | **310** | 342 | 333 | 330 |
|  | *AIC* | 376 | 317 | 305 | **292** | 335 | 319 | 312 |
|  | *Var* | 0.225 | 0.176 | 0.165 | **0.155** | 0.197 | 0.174 | 0.168 |

Non-linear probability weighting is a further subjective factor explaining economic choices, as proposed in prospect theory and other generalized EU theories (Harless and Camerer, 1994; Hey and Orme, 1994). We found that a model incorporating utility and weighted probabilities (PW model) improved the description of the measured IPs by 4.5 times in Monkey A and 4.4 times in Monkey B, compared to the EV model. Therefore, the PW model had similar descriptive power compared to the EU model. Overall, the PW model outperformed the EU model in 3 out of 4 model-accuracy metrics, in both monkeys (Fig. 7, Table 6-1), suggesting that in the tested choice situation adding the subjective weighting of probabilities marginally improved the description of preferences compared to EUT, representing a possible refinement to our EU model for describing preferences (Fig. 6c).

**Figure 7.**
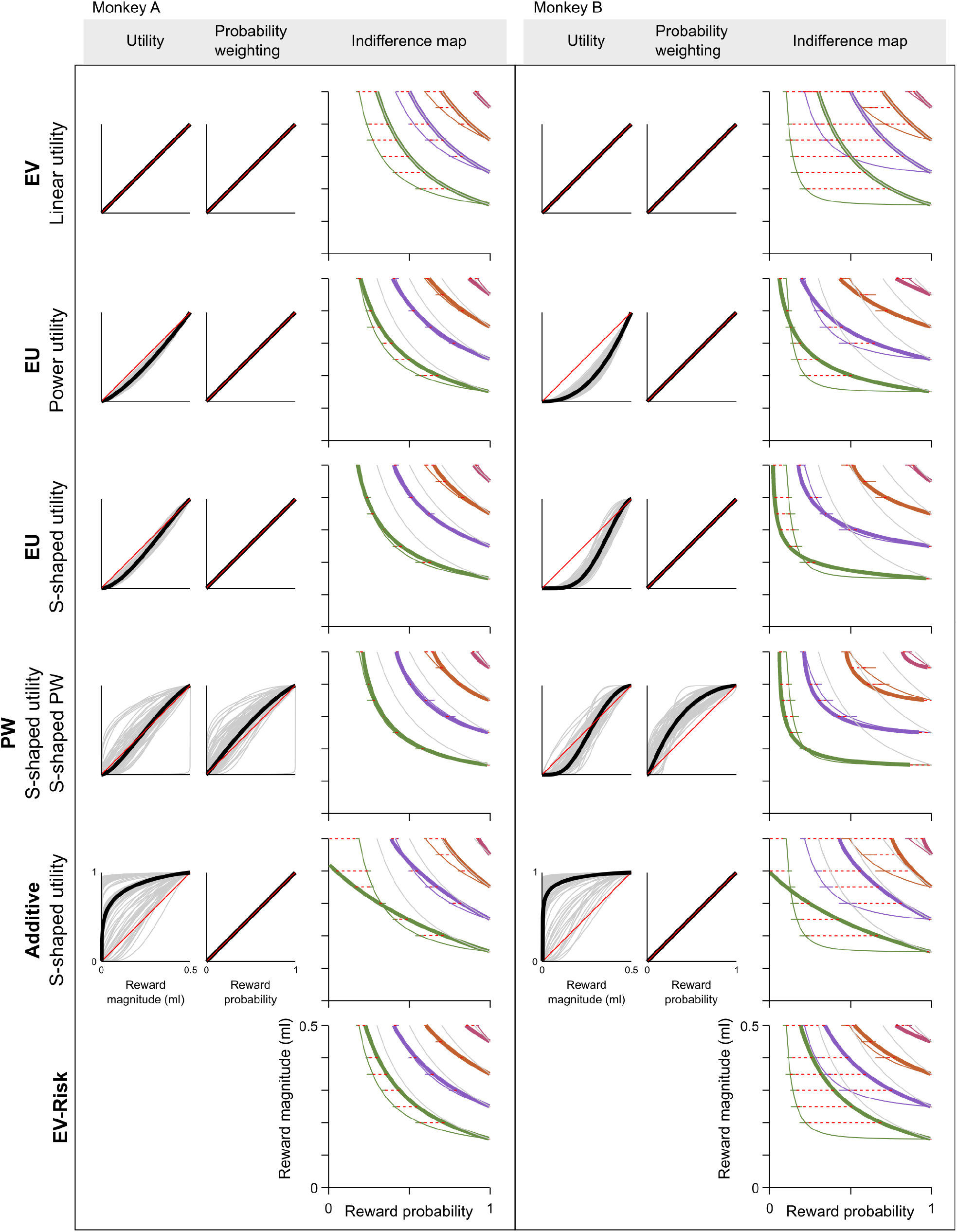
Comparison of economic models. Recovered utility function, probability weighting function and corresponding indifference map for each economic model (rows). Grey curves represent single session estimates, black curves the corresponding means, plotted by averaging the recovered parameters across all sessions. Red lines represent linear utility and probability weighting (PW) functions, for comparison. The EV, EU and additive models assume linear probability weighting. The mean-variance model (EV-Risk) does not have a utility representation. Other conventions and symbols as in Fig. 6b. For numberical comparisons, see Table 6-1.

We explored the possibility of an additive model being able to represent the animals’ choice behavior. Additive models, in which magnitude and probability information are normalized and added to each other (Methods section: *Economic choice models*), have been shown to account for choices under uncertainty (i.e. when risk is not explicitly known) in both humans and monkeys (Farashahi et al., 2019). Fitted to our data, the additive model was outperformed by the other utility-based models (Fig. 7, Table 6-1), confirming the finding that risky choices are better explained by a multiplicative model rather than by an additive one.

The mean-variance approach, an alternative economic model which approximates EUT without relying on the concept of utility (Levy and Markowitz, 1979), defines a gamble’s value as the sum of the corresponding EV and risk components (Methods section: *Economic choice models*). When fitting a mean-variance model to our data, the ICs could not be predicted as well as with any of the utility-based models (Fig. 7, Table 6-1), with an improvement in the ICs description over the EV model of 1.6 (Monkey A) and 1.4 (Monkey B) times, well below the performance of utility-based models.

In support for the existence of a utility-compatible mechanism producing the indifference map, we investigated the variation of IPs across sessions. We computed the Pearson’s correlation coefficient (ρ) for all possible pairs of IPs. A significant ρ, both within each IC (one-sample *t* test, per animal: *p*=1.8·10^−5^ (Monkey A); *p*=5.4·10^−4^ (Monkey B)) and across different ICs (*p*=8.2·10^−5^ (Monkey A); *p*=1.5·10^−3^ (Monkey B)), confirmed that the variation of each IP was associated with a variation of other IPs (average ρ, per animal: 0.19±0.28 SD (A); 0.15±0.26 SD (B)). Across sessions, the indifference map changed shape as a whole: IPs were not varying independently from each other, but were linked by a common underlying root, identifiable as the utility function.

In conclusion, the economic modelling of ICs and the correlation among IPs support the idea of choices resulting from a utility maximization process: the product of a subjectively defined utility function with reward probabilities (possibly subjectively weighted) is able to describe the choice behavior between any pair of gambles and in particular the smooth trade-off relation between reward magnitudes and probabilities (Fig. 8).

**Figure 8.**
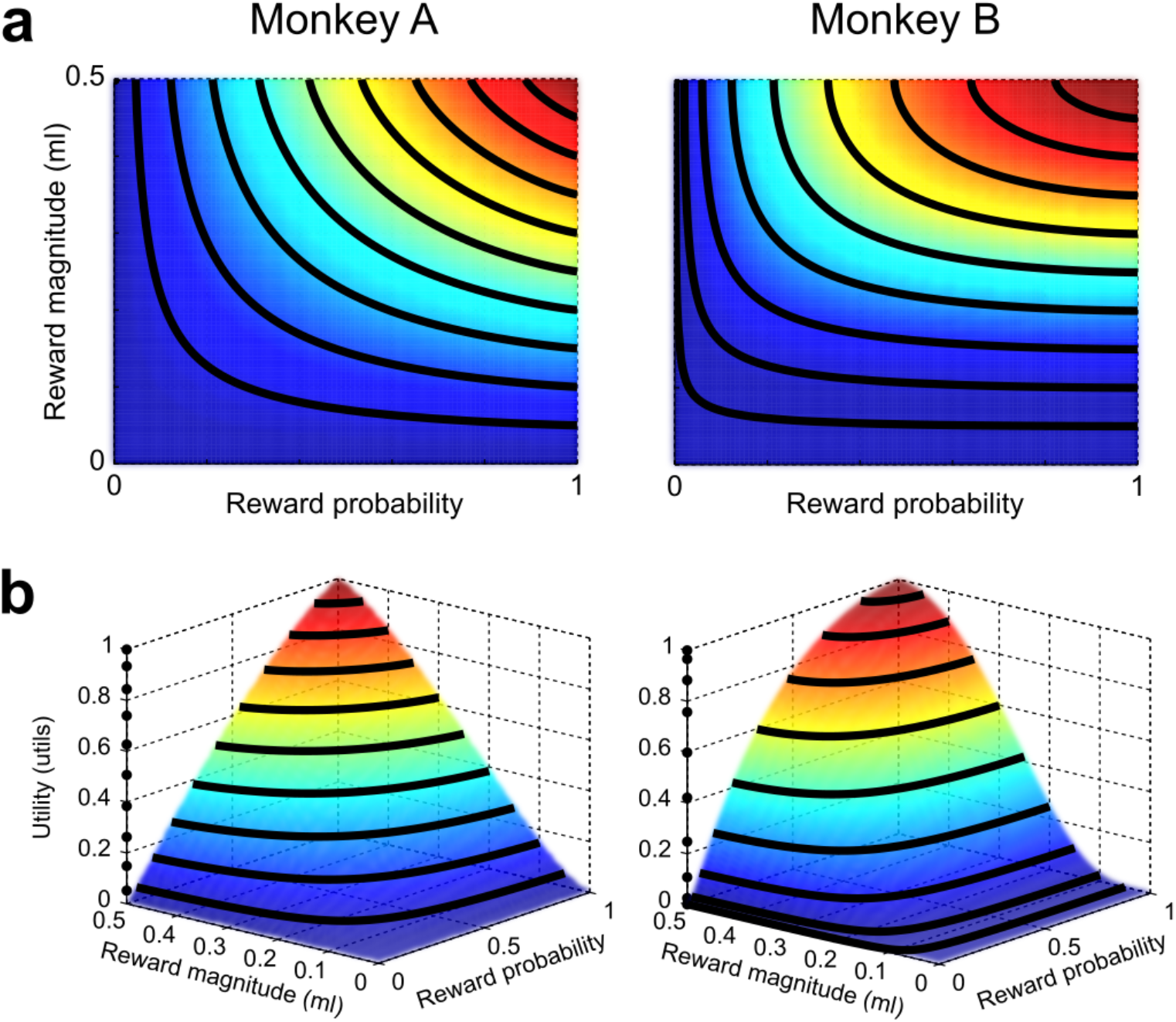
Three-dimensional utility representation. **a) Indifference map resulting from the best-fitting probability weighting model.** Black curves identify the iso-utility points (i.e. the ICs) for 9 equally spaced EV levels (0.05-0.45 ml, step: 0.05 ml). Colors (blue to red) correspond to utility values (from 0 to 1). **b) Three-dimensional representation of utility values,** highlighting the continuous relation between objective quantities (m, p) and subjective values (utility): for any two-outcome gamble, corresponding to a {m, p} pair, a utility level can be mathematically computed using the two-dimensional function defined by the best-fitting PW model (V(m,p) = U(m) · w(p)). This relation can be used to identify utility-coding neurons: the activity of a neuron coding utility should follow the three-dimensional surface across the whole tested region of the MP space.

### Continuity axiom test in the Marschak-Machina triangle

The Marschak-Machina triangle (Marschak, 1950; Machina, 1982) has been extensively used in economic studies of human behavior for evaluating and comparing different generalized EU theories (Camerer, 1989; Sopher and Gigliotti, 1993). This approach graphically displays continuity tests by showing IPs in choices between test gamble B and probabilistic AC combinations containing multiple possible outcomes.

We further tested the continuity axiom using A, B and C gambles defined as two-outcome gambles, which resulted in AC combinations being three-outcome gambles. To present such gambles to the animal, we used visual cues with three horizontal lines, which simultaneously represented all possible reward outcomes and their probabilities (Fig. 9a, inset). The Marschak-Machina triangle represents gambles with three fixed outcome magnitudes (defined in our experiment as 0 ml, 0.25 ml and 0.5 ml) and any combination of associated probabilities (p_1_, p_2_ and p_3_, defined as the probabilities associated with the low, middle and high outcome magnitudes, respectively). The x and y coordinates correspond to the probability of obtaining the lowest (p_1_) and highest (p_3_) outcome, respectively (Fig. 9a).

**Figure 9.**
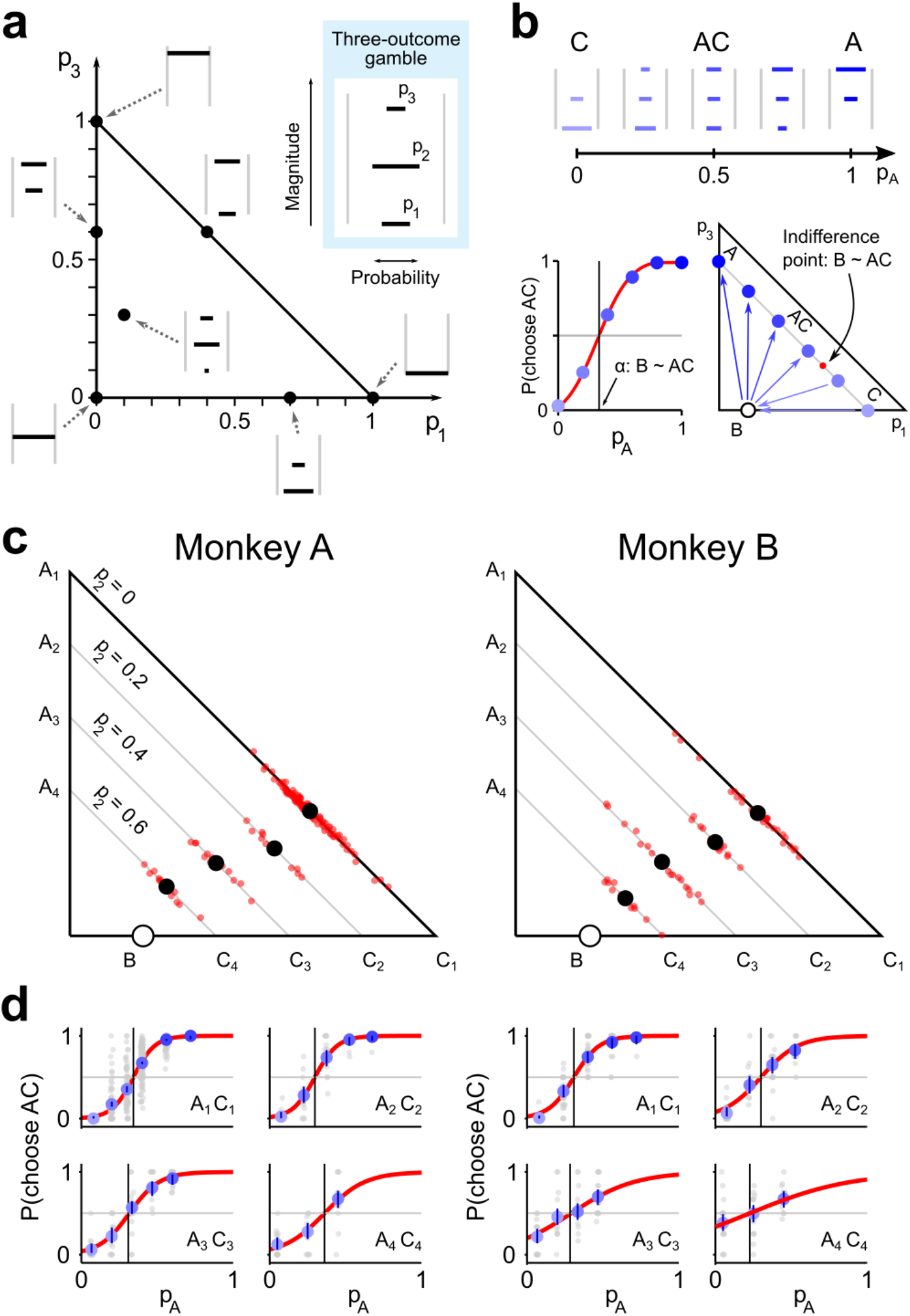
Continuity axiom test in the Marschak-Machina triangle. **a**) **Three-outcome gambles**. Gambles with three fixed outcome magnitudes and any combination of outcome probabilities can be represented in the Marschak-Machina triangle. The visual cue (inset) for three-outcome gambles included three horizonal lines representing the three possible outcome magnitudes (vertical position) and the respective probabilities p_1_, p_2_ and p_3_ (line width). **b**) **Scheme of the continuity axiom test.** Three-outcome gambles used to test the axiom (top) can be represented in the Marschak-Machina triangle (bottom right) together with the B gamble (circle) and the resulting IP (red dot). The arrows point toward the preferred option, consistently with the proportion of AC choices (bottom left). **c) Continuity axiom test in the Marschak-Machina triangle.** Average IPs (black dots) and from single sessions (red dots) were consistently elicited, indicating compliance with the continuity axiom in choices between two- and three-outcome gambles. **d) Revealed preferences in choices between two- and three-outcome gambles.** Average measured percentage of AC choices as a function of the probability of obtaining the A option (graded blue dots). Each average IP (black vertical line) corresponds to a black dot in panel **c**. Other symbols as in Fig. 3.

We defined A as a gamble with 0.25 ml and 0.5 ml as possible outcomes, whereas both B and C had 0 ml and 0.25 ml as possible outcomes; AC combinations then corresponded to gambles with three possible reward magnitudes: 0 ml, 0.25 ml and 0.5 ml (Fig. 9b). In the Marschak-Machina triangle, gamble A lay on the y axis while gambles B and C lay on the x axis. Consequently, the AC(p_A_) combinations lay inside the triangle, on a straight line between A and C, the position between bottom right and top left being proportional to the probability pA (Fig. 9b, bottom). Satisfaction of the continuity axiom would be manifested as a point on the line between A and C where the animal is indifferent between the B gamble and the AC combination (labeled B~AC in Fig. 9b, bottom).

We defined four pairs of A and C gambles (A_1_C_1_ to A_4_C_4_, associated with increasing probability of the middle outcome (p_2_) between p_2_ = 0 and p_2_ = 0.6, in 0.2 increments); for each A-C pair we tested the continuity axiom using a fixed middle gamble B, for a total of four tests (Fig. 9c). Results showed the existence of IPs in all tested cases (Fig. 9d), confirming compliance with the continuity axiom in choices between two- and three-outcome gambles. Because reward magnitudes are fixed in the Marschak-Machina triangle, while probabilities vary across the full range, the pattern of IPs confirmed the role of reward probabilities as modifiers for the EU: a gradual change in A-C (in terms of p_2_) lead to a continuous increase in IPs (in terms of the probability of the highest outcome, p_3_), also demonstrating the possibility of constructing ICs within the Marschak-Machina triangle (Fig. 9c).

Through the unique graphical representation of the Marschak-Machina triangle, we showed that monkeys complied with the continuity axiom in choices involving three-outcome gambles, thus supporting the idea of a choice mechanism based on numerical subjective values also in more complex choice scenarios.

## DISCUSSION

This study demonstrates compliance of monkey behavior with the continuity axiom of EUT, implying a magnitude-probability trade-off relation and determining a numerical utility measure able to describe choices. Our results build on studies reporting monkeys’ preferences for gambles with larger EV to those with smaller EVs derived from both reward magnitude and probability (Musallam et al., 2004; Lau and Glimcher, 2005; Averbeck, 2015; Farashahi et al., 2018). The interpretation of an apparent trade-off between reward magnitude and probability in these studies relies on the validity of EUT axioms in general and on the continuity axiom in particular. By showing compliance with the continuity axiom, we demonstrate that the trade-off is systematic and quantitatively complies with theoretically ideal choices. In addition, the compliance with the continuity axiom fulfils a necessary condition for the validity of neuronal coding of subjective value and formal economic utility (Platt and Glimcher, 1999; Tremblay and Schultz, 1999; Padoa-Schioppa and Assad, 2006; Kobayashi and Schultz, 2008; Lak et al., 2014; Stauffer et al., 2014). These results demonstrate compliance with the continuity axiom for biologically plausible reward magnitudes and probabilities. The animals’ trade-off choices followed closely the IPs modelled by theoretical choice functions, suggesting the existence of an axiomatically defined utility measure for choice options’ values. The pattern of subjective IPs allowed us to define a utility function that is suitable for investigating neuronal mechanisms of economic choice according to the rigorous definitions of economic theory.

The continuity axiom, a necessary condition for the existence of a numerical utility, states that given three subjectively ordered gambles, a decision maker will be indifferent between the middle gamble and a probabilistic combination of the two other gambles. We experimentally tested the continuity axiom in choices between a two-outcome gamble and a safe option. Four monkeys exhibited a choice behavior consistent with the continuity axiom, making choices compatible with the existence of a unique IP. We generalized our results to more complex choice situations in two monkeys, confirming compliance with the axiom in choices between two- and three-outcome gambles, representable in the Marschak-Machina triangle. We showed how the IPs identified through the axiom test procedure could be interpreted as subjective evaluations of the choice options and used to construct an indifference map. Such a map revealed a congruent, subjective trade-off relation between reward magnitudes and probabilities, which supported the idea of choices being the result of a utility maximization process compatible with EUT.

The four axioms of EUT represent the necessary conditions for the existence of a precisely defined utility quantity. In particular, the continuity axiom permits the definition of a numerical utility, while the independence axiom defines how to compute the utility measure. In our quest for investigating a utility-based brain mechanism driving human decisions, we need to clarify if and to what degree the economic theories are generalizable across primates. Although the continuity axiom has not been tested in human subjects, it is accepted as a reasonable condition. On the other hand, humans have been shown to violate the independence axiom of EUT, which led to the creation of alternative economic choice theories. Though it is still unknown whether non-human primates violate the independence axiom similarly to humans, as a first step we showed that they comply with a more basic assumption, the continuity axiom. By sequentially testing the EUT axioms, we can verify up to which point their behavior can be described by the economic theory, and if monkeys’ preferences reflect the characteristics of human decision making. This approach can shed light on the existence of a common choice mechanism across primates.

Past studies have shown that decisions in monkeys reflect both the magnitude and probability information of the choice options (Platt and Glimcher, 1999; Lak et al., 2014), leaving two open questions: how are magnitude and probability, two physical quantities, transformed into subjective quantities? And how are such quantities combined into a single value? These questions naturally extend to the neurophysiological domain. The axiomatic approach allows to investigate such points, clarifying with a robust procedure if the gambles’ dimensions are subjectively combined as mathematically defined by modern economic decision theories.

Lexicographic preferences and other classes of choice heuristics represent known continuity axiom violations. By showing compliance with the continuity axiom we could exclude an important class of heuristics (the lexicographic rules) as the driving mechanism for choices in the tested situation. This ensured that all presented information were used to make actual multi-attribute choices: reward probability and magnitude were both considered and combined when evaluating the options. However, we did not test other heuristic decision strategies. For example, a recent study observed a win-stay/lose-switch strategy, which only contributed marginally to single-trial choices while possibly influencing the long-term learning of values and probabilities (Ferrari-Toniolo et al., 2019). Thus, further tests should delimit the viability of continuity satisfaction in different choice situations, while exploring alternative heuristic-based models able to describe the observed choice patterns.

An important consequence of continuity axiom violations is that subjective preferences cannot be described by assigning numerical values to the options. IPs, and hence indifference maps, would be undetermined if the axiom is violated, as would the concept of utility as a subjective measure of an option’s value. An EUT model could still be fitted to the data to recover a utility function; yet, such a function would not have the intended meaning of expressing the options’ subjective values. When the axiom is fulfilled, instead, we showed how the IPs could be expressed numerically as utility, and the resulting indifference map could be generated through s-shaped utility functions. Having a utility representation of values allows for the assignment of a specific numerical value to each indifference curve. The activity of a neuron encoding the options’ subjective values should comply with the indifference map: it should be proportional to the elicited numerical utility levels across indifference curves, while remaining constant within each indifference curve (see Fig. 8).

The utility function is not considered to be fixed over time: its shape and range has been theorized to adapt and to change based on internal and external factors. Multiple factors might contribute to the shaping of each subject’s utility function in different ways. Some of the observed variability in our utility measure might be explained as an adaptation to changes in the task or in the external environment, as suggested by the gradual changes observed in IPs across sessions. This calls for more sophisticated models integrating longer term learning effects on the formation and the evolution of utility functions with experience.

According to theories relying on the continuity axiom, utilities are combined with probability information to give a gamble’s EU. The exact form of such combination remains to be tested: the independence axiom is required to define exactly how utilities and probabilities combine into EU. Although we did not yet explicitly test compliance with the independence axiom in monkeys, we observed that non-linear weighting of reward probabilities resulted in a marginally better description of choice behavior compared to EUT. Therefore, the present study points to non-linear probability weighting as a possible refinement to EUT in monkeys, compatible with the human experimental results that led to the development of prospect theory (Kahneman and Tversky, 1979). The Marschak-Machina triangle framework could be used to directly investigate compliance with the independence axiom in monkeys, allowing for the quantitative investigation of the neural underpinnings of several generalized EU choice theories.

In conclusion, by explicitly testing the continuity axiom we verified that, in the tested situation, the choice mechanism was compatible with the computation of finite, numerical utilities, gaining crucial information on the plausible mechanisms guiding choices toward the maximization of utility.

## Acknowledgments

This work was supported by Wellcome Grants WT 095495 and WT 204811 and European Research Council Advanced Grant 293549. We thank Aled David and Christina Thompson for animal and technical support, and Federico M. Echenique and David M. Grether (Caltech) for helpful comments.

